# All signals considered: Data quality partially explains inter-individual task differences in a large, open fNIRS dataset

**DOI:** 10.64898/2026.06.06.728412

**Authors:** Sophie Raible, João Pereira, Foivos Kotsogiannis, Bruno Direito, Teresa Sousa, Manuela da Cunha Seiffert, Rik Lavicka, Vendija Skeltona, Daniëlle Evenblij, Assunta Ciarlo, Armin Heinecke, Jacqueline Gädtke, Zeus Tipado, David M. A. Mehler, Simon H. Kohl, Miguel Castelo-Branco, Rainer Goebel, Michael Lührs, Bettina Sorger

**Author notes:** The first two authors contributed equally to this work.

## Abstract

**Significance:** High inter-subject variability and limited reproducibility in functional near-infrared spectroscopy (fNIRS) research may partly reflect global systemic physiology and signal quality differences, possibly distorting task-evoked hemodynamic responses.

**Aim:** We investigate how signal quality relates to inter-subject variability in motor-task fNIRS responses and introduce a large, open, multi-task, near whole-head fNIRS dataset with extensive peripheral physiology and short-channel recordings.

**Approach:** Fifty-seven participants completed resting-state, motor action, motor imagery, emotion recognition, visual, and auditory tasks during fNIRS recording. Peripheral measures included pulse oximetry, heart rate, blood oxygen saturation, respiration, room temperature, galvanic skin response, electrocardiogram, and electromyography. Signal quality was assessed using the scalp coupling index (SCI), coefficient of variation (CV), signal-to-noise ratio (SNR) and a spectral measure here coined the coupling SNR (cSNR).

**Results:** Quality metrics were weakly to moderately correlated, except SNR and CV, which showed the expected inverse relationship. All quality metrics were significantly related to channel length and associated with task-related activation estimates. Group-level analyses validated activation in expected task-related regions.

**Conclusions:** The assessed metrics capture complementary features of fNIRS signal quality and may help explain individual activation differences. The dataset provides a comprehensive, open resource enabling future evaluation of physiological correction methods and confound mitigation.

## 1 Introduction

Functional near-infrared spectroscopy (fNIRS) is an optical brain imaging method using near-infrared light (700-900 nm) to measure local changes in light absorption, allowing for an indirect quantification of neuronal activity through the estimation of oxygenated hemoglobin (HbO), deoxygenated hemoglobin (HbR) and cerebral total hemoglobin (HbT).^1–3^ fNIRS allows for the noninvasive investigation of brain activity in naturalistic task settings and has gained increasing popularity in the last years.^4,5^

The portable and relatively movement-tolerant nature of fNIRS makes it an attractive tool for personalized applications such as rehabilitation monitoring,^6^ neurofeedback interventions^7,8^ or brain-computer interfaces.^9–13^ However, while task-related effects are often evident at the group level, fNIRS responses show substantial inter-subject variability,^14–17^ where some participants show robust, spatially plausible hemodynamic activation, whereas others show weak, absent, atypical, or noisy responses.^14^ Although these differences are often interpreted as reflecting meaningful activation changes, they might also be explained by systemic physiological influences,^18–20^ extracerebral hemodynamics,^18–20^ or differences in measurement quality caused by optical coupling, detector saturation or motion artifacts. Since individual activation changes are especially important in personalized applications, understanding where differences in task-evoked fNIRS activity stem from is especially important for the future development of interventions such as fNIRS-based neurofeedback therapy or brain-computer interfaces for motor-independent communication and control.

Several approaches have been proposed to quantify fNIRS signal quality. At the raw-signal level, channels are commonly assessed using metrics such as the signal-to-noise ratio (SNR), coefficient of variation (CV) or detector saturation.^21,22^ Because well-coupled fNIRS channels should contain physiological oscillations (caused by the heart beat), several quality metrics also evaluate the presence of cardiac pulsatility, including the scalp coupling index (SCI)^23,24^ and related spectral measures.^25^ More recent approaches combine multiple signal features into automated or semi-automated quality indices, such as the signal quality index (SQI), which rates fNIRS signal quality on an ordinal scale.^26^ Overall, there is no single universally accepted metric for fNIRS signal quality. Current best practice recommendations therefore emphasize that any criteria used should be reported explicitly.^22^ While the field has made an effort towards reporting these metrics, signal quality is mostly treated as a nuisance factor: Poor channels are removed, data are filtered, and analyses proceed on the remaining signals. Much less is known about whether signal quality might have a confounding effect on typically reported task-related activation changes.

In addition to signal quality measures, awareness of physiological influences on the fNIRS signal has increased in the field over the last years. Since the near-infrared light measured by fNIRS travels through extracerebral tissue layers such as the scalp before reaching the brain, it is strongly influenced by local changes in blood flow and oxygenation in these superficial layers, which can mask or exaggerate neurally driven hemodynamic responses.^18,27^ Several instrumentation-based^17,20,28–31^ or data-driven approaches^32^ exist for dealing with physiological confounds in the fNIRS signal. While existing approaches have substantially improved the correction of systemic physiological contributions to the fNIRS signal, rigorous quantitative evaluation remains challenging because suitable public datasets are limited. As a result, both established and emerging methods for controlling physiological influences have largely been validated using small, task-specific datasets^31,33^ or simulation approaches.^29,34–36^ The lack of large-scale datasets that explicitly capture physiology makes it difficult to determine best practices for physiology correction.

Here, we introduce and validate a comprehensive, large, openly available fNIRS dataset of n = 57 subjects, with near whole-head fNIRS, full short-channel coverage, and comprehensive physiological recordings from multiple task domains (including resting state, motor action and imagery, emotion recognition, auditory, and visual tasks). Beyond empirical investigations of data quality contributions to inter-individual differences in task-evoked fNIRS responses, this dataset also facilitates the systematic evaluation and development of physiology-correction methods, thereby informing (task-dependent) recommendations and best practices. As an initial application, we report a broad set of signal quality metrics and test whether variability in these measures can partially explain inter-individual differences in activation patterns during a motor paradigm.

## 2 Materials and Methods

### 2.1 Participants

A total of n = 57 subjects (mean age: 26.93 ± 8.67 years, 59.65% female) participated in this study. Participants had no neurological or psychiatric disorders and normal or corrected-to-normal vision. Written informed consent was obtained from all participants before data collection. The study was conducted in accordance with the Declaration of Helsinki and approved by the Ethics Review Committee Psychology and Neuroscience of Maastricht University (ERCPN-255_99_07_2022). Sample size was determined pragmatically by the available time and resources, and data collection aimed to obtain at least 20 datasets with acceptable fNIRS signal quality, complete task and questionnaire data, and usable physiological recordings. This target was chosen a priori for practical reasons rather than based on a formal statistical justification or power calculation.^37^

### 2.2 fNIRS measurements

fNIRS data were recorded using Aurora v2023.4 with two cascaded NIRSport2 continuous-wave (CW) fNIRS systems (Nirx Medical Technologies, Berlin), using 32 light-emitting diode (LED) sources (λ_1_ = 760 nm; λ_2_ = 850 nm) and 28 Silicon Photodiode (SiPD) detectors sampled at 12.6 Hz. Each source was equipped with one of 32 short-distance detectors (SDD), placed ca. 8 mm from the respective source, resulting in 102 regular and 32 short-separation channels (total 134 channels). Head movements were recorded using a 9-axes (3 linear acceleration, 3 gyroscope, 3 magnetometer) accelerometer integrated in the used fNIRS system. Functional coverage was designed using NIRSite v2021.4 and the fNIRS Optodes Location Decider (fOLD) toolbox.^38^ The resulting layout, displayed in Fig. 2 (a), was confined to the 10/10 system and aimed at providing comprehensive cortical coverage, focusing especially on occipital (primary visual cortex - V1) and motor-related cortices (bilateral primary sensory and motor cortices - S1 and M1, premotor cortices - PMC, and supplementary motor areas - SMA), the temporoparietal junction (TPJ), prefrontal (dorsolateral prefrontal cortex - dlPFC), and auditory areas. Source and detector positions were projected onto the scalp surface in Satori 2026.4, and wavelength-specific sensitivity Jacobians were estimated on the gray-matter mesh using a semi-infinite slab model. The resulting channel sensitivities were summed over the montage and log10-transformed to generate the sensitivity profile displayed in Fig. 2 (b).

**Fig. 1.**
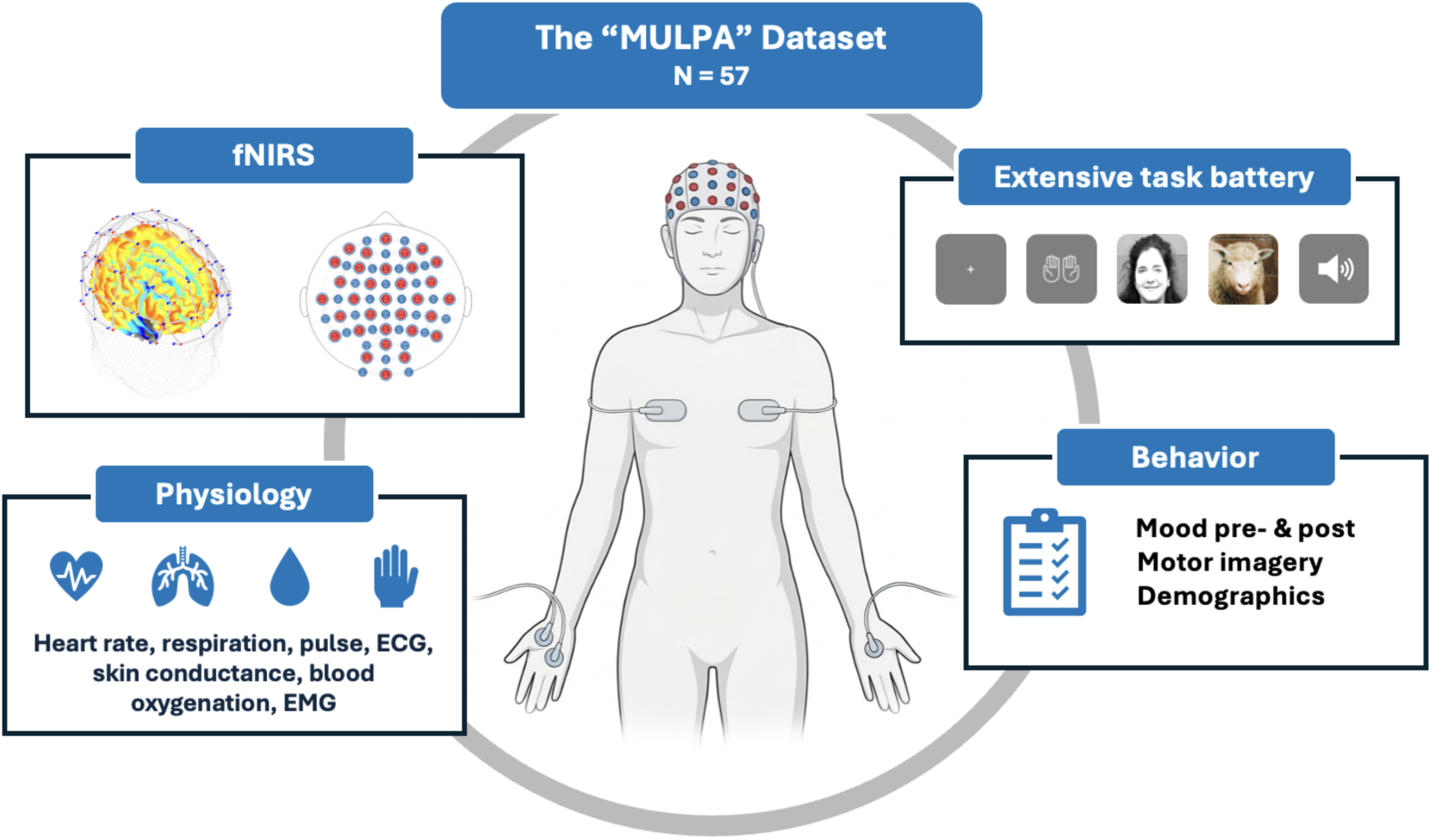
Overview of the entire BIDS-compliant “MULtiple PAradigms (MULPA)” dataset, including fNIRS montage and sensitivity profile. Physiology measures include heart rate, respiration, pulse (PPG), an electrocardiogram (ECG), skin conductance (GSR), blood oxygenation (SpO2), and electromyography (EMG) at the hand. Tasks include resting state, motor, emotional, visual, and auditory tasks.

**Fig. 2.**
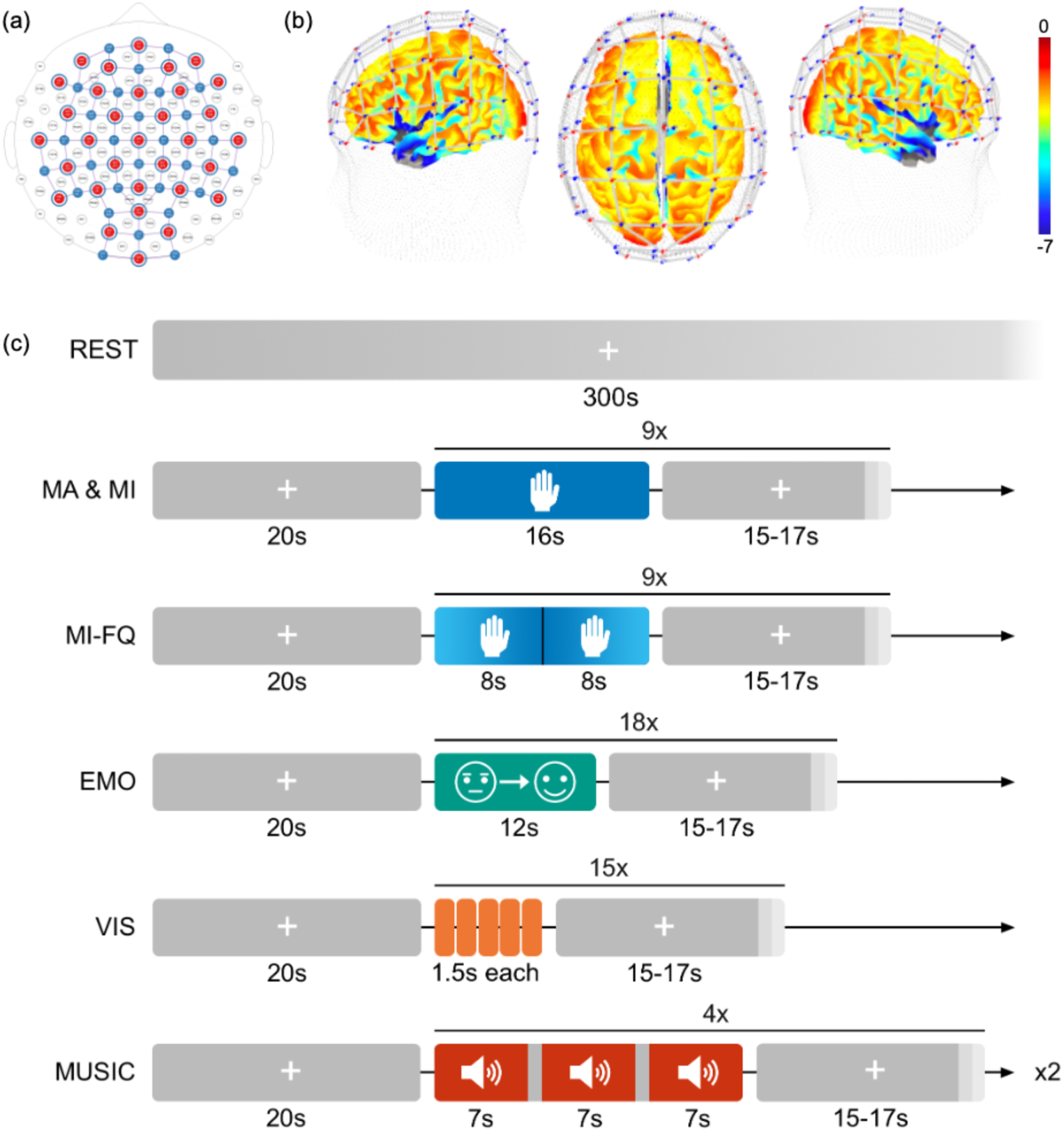
(a) fNIRS montage with 32 sources, 28 detectors and 32 short-distance detectors. (b) Sensitivity profile in left-hemispheric (left), top (middle) and right-hemispheric (right) view. Sensitivity values are expressed as log10 scale. (c) Overview of all task timings. Gray blocks represent rest periods, colored ones task periods. REST = Resting State, MA = Motor Action, MI = Motor Imagery, MI-FQ = Motor Imagery with frequency change, EMO = Emotion, VIS = Visual.

### 2.3 Physiological measurements

External physiological signal measurements were performed using the NIRx Wings 1 module (Nirx Medical Technologies, Berlin). Global body parameters recorded included pulse oximetry (PPG), heart rate (HR), oxygen saturation (SpO2), respiration, galvanic skin response (GSR), and an electrocardiogram (ECG). Additionally, electromyography (EMG) was measured at the right hand to detect any muscle activity and monitor for subtle movements, particularly important in the context of motor imagery, and room temperature was captured. The NIRx Wings module was coupled with NIRSport 2 through the Aurora software via Bluetooth, and all signals were fully synchronized in one data Lab Stream Layer (LSL) stream.

### 2.4 Experimental protocol

At the start of the measurement, participants completed the Motor Imagery Questionnaire (MIQ3)^39^ and the original Profile of Mood States (POMS).^40^ The MIQ3 is a self-assessment of the ability to perform covert motor tasks. The POMS is a psychological rating scale used to assess transient, distinct mood states. After completing the questionnaires, the fNIRS cap was placed on participants’ heads, ensuring an equal distance of Cz to nasion and inion, and Cz to the two preauricular points. fNIRS and physiological measurements were conducted for an extensive task battery consisting of five distinct parts: (1) A resting-state measurement, (2) three different motor tasks, (3) an emotion-recognition task, (4) a visual task, and (5) a music-listening task. An overview of the task protocols is shown in Fig. 2. The order of tasks was fixed across participants. All tasks started with a 20-seconds baseline block. Task blocks were always interleaved with baseline periods of variable length ranging from 15 to 17 seconds in 1-s increments, during which a fixation cross was presented. After completion of the task battery, participants completed the post-measure of the POMS questionnaire. Physical characteristics such as skin and hair color were assessed with an in-house developed experimenter-rated questionnaire.

#### 2.4.1 Resting-state

During the resting-state task, a fixation cross was presented on the screen for five minutes. Participants were instructed to relax, focus on the cross, and not think of anything in particular. The relatively short resting-state duration was chosen as studies suggest that functional connectivity estimates can be obtained even from short acquisitions in fNIRS.^41,42^

#### 2.4.2 Motor tasks

Motor tasks consisted of 9 blocks of 16 seconds, in which an icon displaying two hands instructed participants to perform the task. In the motor action task, participants were instructed to perform bilateral finger tapping at a consistent frequency. In the motor imagery task, subjects were instructed to imagine a bimanual movement at a consistent speed. While participants were asked to try to imagine the same finger-tapping movement as in the motor action task, they were allowed to switch to other bimanual movements, such as raising and lowering their arms, if they had difficulty imagining this movement. In the third motor task, participants were instructed to perform a bimanual motor-imagery task with a within-block gradual increase, followed by a gradual decrease in the frequency of the imagined movement.^43^ The switch was indicated to participants with an auditory stimulus at the middle time point (after 8 seconds) of the task block. In total, each motor task run lasted approximately 5.5 minutes.

#### 2.4.3 Emotion recognition task

In the emotion recognition task, participants were presented with dynamic emotional transitions in human facial expressions. The facial expressions in these pictures always started as neutral and then slowly morphed into either a sad (Neutral-Sad condition) or happy expression (Neutral-Happy condition), or remained neutral (Neutral-Neutral condition). Images were extracted from the Extended Cohn-Kanade Dataset^44^ and serve as representative keyframes of the dynamic stimuli used. The order of conditions was randomized. A total of 18 blocks (six for each condition) comprised the experimental run, with each block lasting 12 seconds. In total, the run lasted approximately 8 minutes.

#### 2.4.4 Visual task

In the visual task, participants were presented with images of threatening or non-threatening animal faces, or control stimuli consisting of grass or flowers.^45^ The visual task run consisted of 15 blocks (five per condition) of 7.5 seconds, where five images appeared on the screen for 1.5 seconds each, resulting in an overall run length of approximately 5 minutes.

#### 2.4.5 Music listening task

During the music listening task, participants were asked to close their eyes and listen to music excerpts. The excerpts were selected from a public dataset of 900 30-seconds audio clips.^46^ Based on Russell’s circumplex model of emotion,^47^ we established four experimental conditions by mapping the valence and arousal of the musical pieces into four distinct quadrants: Q1 - Positive Valence, High Arousal; Q2 - Negative Valence, High Arousal; Q3 - Negative Valence, Low Arousal; Q4 - Positive Valence, Low Arousal. The order of the conditions was randomized. A total of 12 excerpts (three for each quadrant) were played during the experimental run, with each excerpt lasting 7 seconds, and a 1-second inter-stimulus interval, resulting in a total run length of approximately 3 minutes. This task was performed twice.

### 2.5 Dataset and availability

The dataset is organized according to the Brain Imaging Data Structure (BIDS) specification for fNIRS (BIDS version 1.11.0) and is openly available in the repository “The MULtiple PAradigms (MULPA) dataset” on Zenodo (https://zenodo.org/records/21033499). Raw continuous recordings are provided in .snirf format (sub-**_task-**_nirs.snirf), which contain the optical intensity measurements as well as synchronized auxiliary physiological signals (PPG, HR, SpO2, respiration, temperature, GSR, ECG and EMG) acquired during the experiment. Each fNIRS recording is accompanied by required BIDS sidecar files, including JSON metadata files describing acquisition parameters and TSV files detailing channel information (_channels.tsv), optode positions (_optodes.tsv), and the coordinate system (_coordsystem.json). As the optode setup was consistent for the whole study, optodes.tsv files are identical across participants. Participants are ordered according to completeness and quality of the data: The first 22 participants constitute full datasets (complete questionnaires, complete data for at least one run of each task) of good data quality (defined as SCI < 0.75 for a maximum of 33% of all channels, and no missing data points or signal drifts in physiological data based on visual inspection). The following 35 participants are ordered in descending quality and completeness of data, an overview of which can be found per participant in the participants.tsv file, as well as in an online form (https://sophieraible.github.io/mulpa-dataset-website/), which allows for filtering by participant and run and simplifies data selection.

The seven distinct fNIRS tasks included in the dataset are labelled task-restingstate, task-motoraction, task-motorimagery, task-motorimageryfreqchange, task-emotion, task-visual and task-music. Task timing and experimental annotations are provided in *_events.tsv files, which specify event onsets, durations and trial types. Task-level metadata shared across participants are listed in corresponding *_nirs.json files.

Pre- and post-experimental questionnaires are stored as TSV files (e.g. poms_pre.tsv) in the phenotype sub-directory, with accompanying JSON sidecars describing questionnaire variable definitions. Participant-level demographic and descriptive information, including data quality and completeness, is provided in participants.tsv and participants.json.

### 2.6 fNIRS data preprocessing

fNIRS data were preprocessed using Satori 2026.4. After conversion of the raw fNIRS data to optical density changes, channels with an SCI of < 0.75 were pruned. Data was then converted to relative HbO and HbR concentration using the modified Beer-Lambert Law. Motion correction was performed on the concentration data by applying a spike removal algorithm (with default values of: Iteration = 10, Lag = 5 seconds, Threshold = 3.5, Influence = 0.5 and monotonic interpolation) and temporal derivative distribution repair (TDDR)^48^ to restore high frequencies. A general linear model (GLM) was used to regress out superficial signal components of the highest-correlated short separation channel from the fNIRS signal. Finally, a high-pass Butterworth filter (0.01 Hz) and low-pass Gaussian smoothing (0.4 Hz) were applied, and the data was z-normalized. The Satori .flow file for preprocessing is included in the public data repository.

Preprocessing of the resting state data differed from other tasks to optimize data for functional connectivity analysis. The resting state preprocessing did not include SCI-based channel rejection. The first and last 20 seconds were removed from resting state time courses to reach steady state.^49–51^ Resting state data was band-pass filtered (Butterworth 0.01–0.1 Hz)^50,52–54^ and linear trends were removed.^51,53^ All other preprocessing steps mirrored those performed for the other tasks.

### 2.7 fNIRS data quality analysis

To characterize fNIRS signal quality, we computed the SCI, CV, and SNR of the raw signal in the motor action run of all n = 57 participants. These measures were chosen for being commonly used for channel rejection, as reported in the literature.^22–24^ The motor action task was chosen as it was expected to show the clearest and most localized task-evoked response in the fNIRS signal, facilitating the analysis signal-quality influences on task effects. For each channel and wavelength, SNR was calculated from the raw signal in decibels as

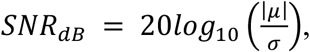

where *μ* and *σ* are the mean and standard deviation of the time-series signal, respectively.^22^ The CV was calculated as the standard deviation divided by the mean raw signal amplitude for each channel and wavelength, and expressed as a percentage:^21^

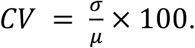

The SCI was calculated in Satori according to Pollonini et al. as the channel-wise zero-lag cross-correlation between the two wavelength-dependent optical density signals after band-pass filtering between 0.5 and 2.5 Hz to isolate the cardiac component and normalizing signal amplitudes, with higher absolute values indicating stronger coupling and thus better signal quality.^23^

To include an additional spectral measure of data quality, we computed a channel-wise metric here termed the “coupling SNR” (cSNR). The aim of this measure was to capture differences in signal quality that could possibly be obstructed in the SCI due to ceiling effects. The cSNR was calculated from the demeaned raw intensity time series for each source-detector pair for each wavelength. Power spectral densities were then estimated independently for the two wavelengths using Welch’s method, with a segment length of 300 samples. Cardiac-related signal power was defined as the integrated spectral power within the frequency range 0.80–2.00 Hz. Noise power was defined as the sum of integrated spectral power within two non-cardiac reference bands, 0.10–0.50 Hz and 2.10–2.40 Hz. Spectral power within each frequency band was obtained by trapezoidal numerical integration of the Welch power spectral density estimate. Cardiac-band power and noise-band power were each averaged across the two wavelengths, and cSNR was expressed in decibels as

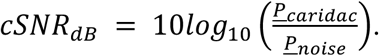

Higher cSNR values indicate stronger cardiac-frequency content relative to off-band spectral power and were interpreted as evidence of better optode-scalp coupling. A detailed overview of how this metric was computed can be found in Appendix Fig. S1.

Associations between channel length and signal quality were assessed using Pearson correlations. Channel length was computed from the .snirf measurement list and 3D probe geometry as the Euclidean distance between source and detector positions. An overview of the channel length calculation process is provided in Appendix Fig. S2. Separate channel-length to signal-quality correlations were computed for SCI, cSNR, SNR, and CV. SNR and CV were averaged across wavelengths for this purpose. To account for multiple testing across the four signal quality metrics, *p*-values were corrected using the Benjamini-Hochberg false discovery rate (FDR) procedure.

To assess whether signal quality was associated with task-related hemodynamic responses, we performed channel-wise ordinary least-squares analysis of covariance (ANCOVA) analyses in Satori using data from all 57 participants. For each participant, the motor action task-effect estimate, representing the task-versus-baseline contrast, was entered as the dependent variable. Signal-quality metrics were evaluated in separate models: for each quality metric, a distinct set of channel-wise ANCOVA models was fitted with that metric as the sole covariate, with no additional confounds included. Benjamini-Hochberg FDR correction was applied separately for each quality metric, chromophore, and model term across the 134 channels. For CV and SNR, HbR and HbO effects were assessed in the metrics derived from the λ = 760 nm and λ = 850 nm time series, respectively. For cSNR and SCI, all the hemoglobin measures (HbO, HbR, and HbT) were assessed.

### 2.8 fNIRS task effects analysis

To assess functional validity of the fNIRS data, we analyzed task effects at the group level. These analyses were restricted to the first n = 22 participants, all of whom met the inclusion criteria of adequate fNIRS data quality, complete physiological sensor recordings, and completion of all experimental runs and questionnaires (as described above). For the resting state task, subject-level functional connectivity was estimated by calculating the Pearson correlation between the time course of each channel pair (excluding short-channels), resulting in a 102 x 102 connectivity matrix per subject. These subject-level matrices were then averaged across participants to generate group-level functional connectivity estimates. To assess across-chromophore consistency, we computed the Pearson correlation between the upper-triangular portions (excluding the diagonal) of the group-averaged HbO and HbR matrices. Statistical significance was evaluated using 100,000 permutations, in which the HbO matrix was randomly shuffled across channels while the HbR matrix was kept fixed. For all other tasks, task-specific group-level random effects (RFX) general linear model (GLM) analyses were run on the HbO data, involving the particular task conditions as GLM predictors. Only one run of the music tasks was included in the analyses per subject.

### 2.9 Physiology analysis

Physiological data were visually assessed, and runs exhibiting clear sensor malfunctions (e.g., signal dropout, saturation, or excessive noise) were excluded from the analysis (respiration: 15/449; heart rate: 46/449, PPG: 6/449, GSR: 26/449, ECG: 10/449, EMG: 7/449 runs). Data quality of the remaining runs was assessed by computing the SNR of each modality. Task differences in mean heart rate and breathing rate were assessed visually. As the aim was to provide an overview of the measurements available in this dataset, no statistical analyses were performed.

## 3 Results

### 3.1 Signal quality of fNIRS data

Figure 3 shows an overview of the data quality across all 57 participants. Panel (a) shows the distribution of SCI, cSNR, SNR, and CV for regular and short fNIRS channels per participant. Panel (b) shows the correlation between these different metrics on the participant-level means, with SNR and CV showing a strong anti-correlation, while the other measures show low to medium correlations. Panel (c) displays the *z*-scored participant means to visualize per-channel (dis-)agreement of the four metrics.

**Fig. 3.**
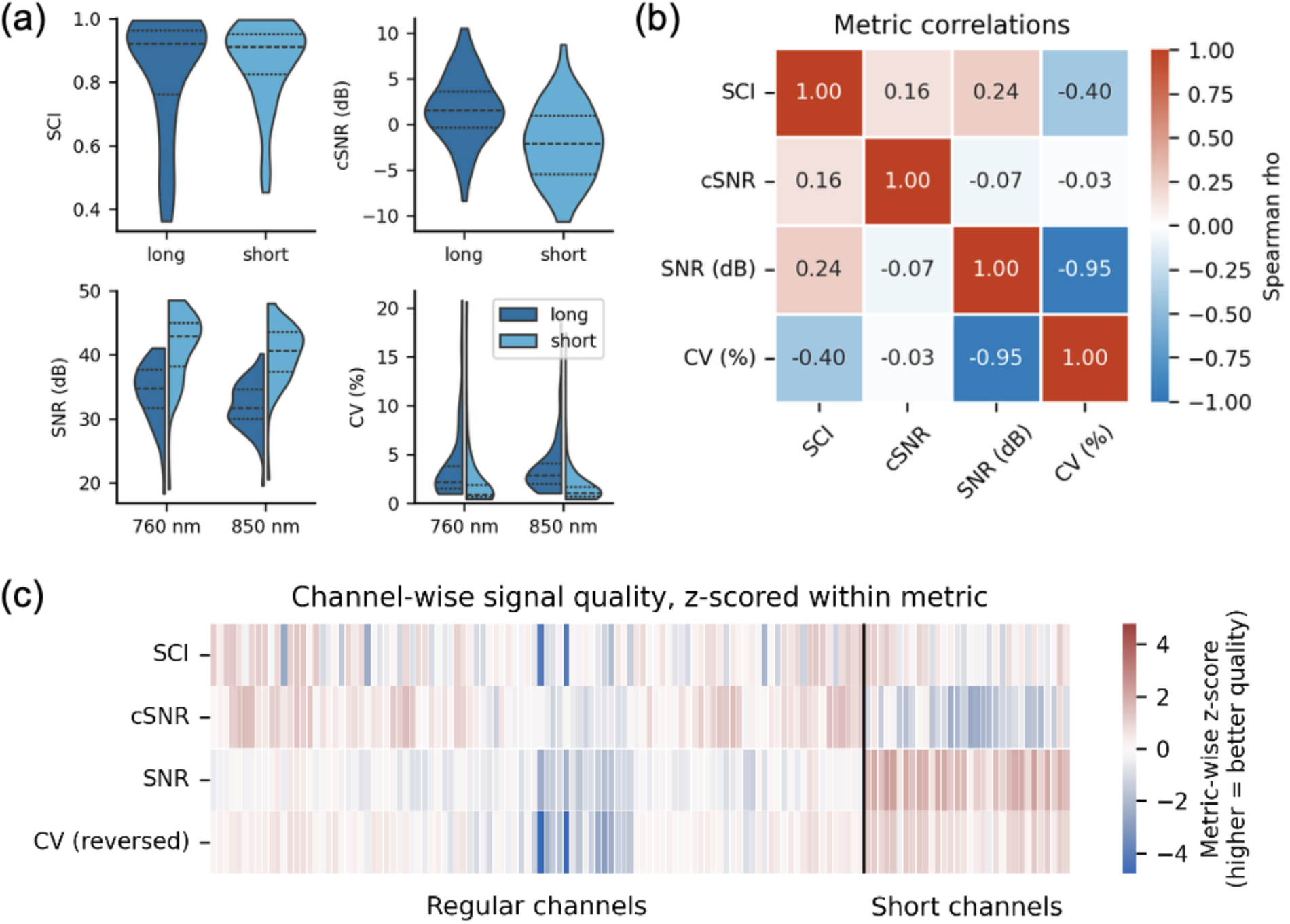
(a) Distribution of signal quality metrics calculated from the motor action task, averaged across channels per subject (n = 57). (b) Correlations between the different signal quality metrics, with SNR and CV values averaged across wavelengths. (c) *Z*-scores of these metrics displayed per channel, with CV values inverted to allow for better comparison. Each column corresponds to one channel. SCI = scalp coupling index, SNR = signal-to-noise-ratio, cSNR = coupling SNR, CV = coefficient of variance.

All four signal quality metrics were significantly associated with channel length after FDR correction. Longer channels were associated with lower SCI (*r* = -0.24, *p*_FDR_ = .006) and lower SNR (*r* = -0.84, *p*_FDR_ < .001). In contrast, channel length was positively associated with cSNR (*r* = 0.67, *p*_FDR_ < .001) and CV (*r* = 0.47, *p*_FDR_ < .001). These results are presented in Fig.4.

**Fig. 4.**
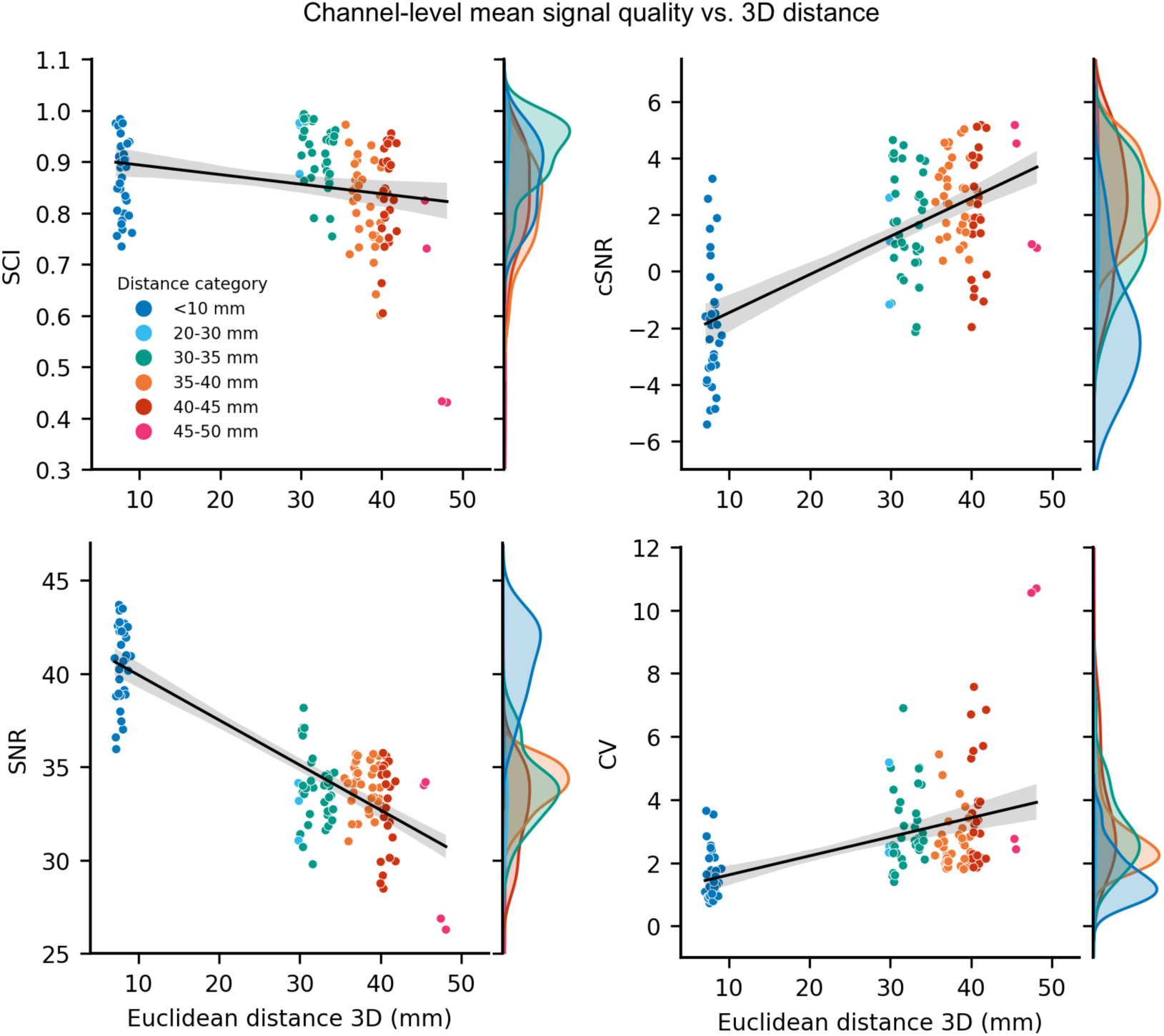
Relationship between channel length (measured as Euclidean 3D source-detector distance) and the channel-level (averaged across participants, n = 57) signal quality metrics in the motor action run. SNR and CV values are averaged across wavelengths. Each point represents one channel, and channels are colored according to distance category. Black lines indicated fitted regression trends for visualization. Side distributions show the marginal distribution of each signal quality metric, split by distance category.

The results of the ANCOVA investigating the effect of signal quality on inter-subject variability in motor task-induced activation are shown in Fig. 5. Each panel displays the *F*-map for a specific quality metric thresholded at FDR-corrected *p* ≤ .05. *F*- and *p*-values for significant channels are reported in Appendix Table S1. Following FDR correction across 134 channels separately for each quality metric, hemoglobin measure, and model term, significant associations were observed for all quality metrics. Higher SCI was associated with more negative HbR task-effect estimates in seven channels and with HbT estimates in two channels, although the HbT effects were directionally mixed. Higher cSNR was associated with task-effect estimates in four HbO channels and one HbR channel, including complementary positive HbO and negative HbR associations in S25-D24. No HbT associations survived FDR correction for cSNR. For the wavelength-specific metrics, higher CV_λ=760_ was associated with more positive HbR estimates in eight channels, higher CV_λ=850_ with more negative HbO estimates in three channels, higher SNR_λ=760_ with more negative HbR estimates in five channels, and higher SNR_λ=850_ with more positive HbO estimates in twelve channels.

**Fig. 5.**
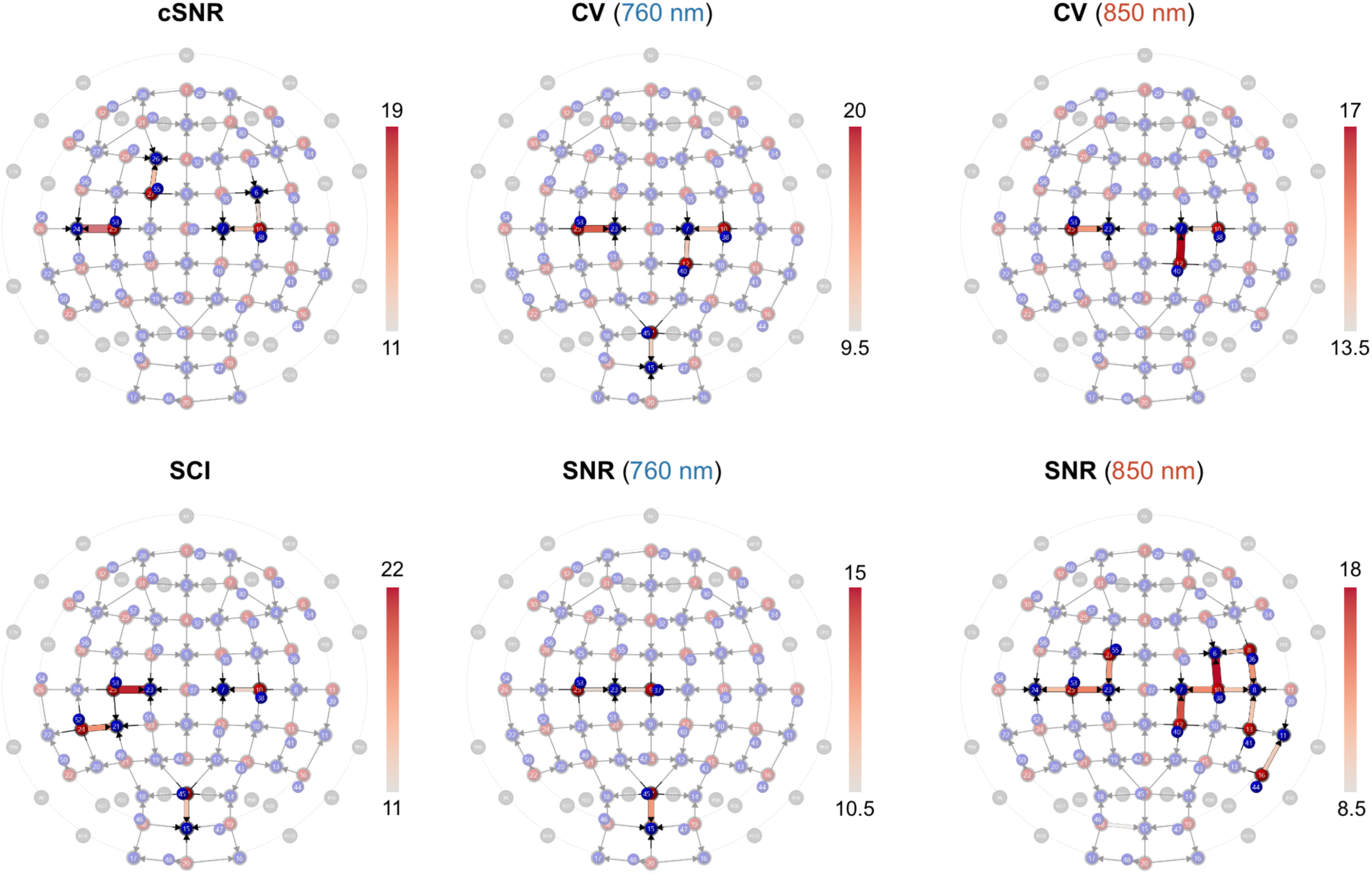
*F*-maps from the ANCOVA, showing the effects of each signal quality metric on task activation in the motor action task across all n = 57 participants. Only channels in which the quality metric covaried significantly (*p* ≤ .05, FDR-corrected) are shown. Significant effects are localized in motor-related channels, suggesting that data quality (partially) explains task-evoked activation in fNIRS signal.

### 3.2 Task effects on fNIRS data

Figure 6 summarizes group-level task-evoked responses. For the resting state data, panel (a) shows the top 5% strongest connections at the group level, illustrating both intra- and inter-hemispheric connectivity. HbO and HbR connectivity matrices revealed a strong correspondence (*r* = 0.82, permuted *p* = 9.9999 × 10⁻⁶), demonstrating cross-chromophore robustness in estimating functional connectivity from this dataset.

**Fig. 6.**
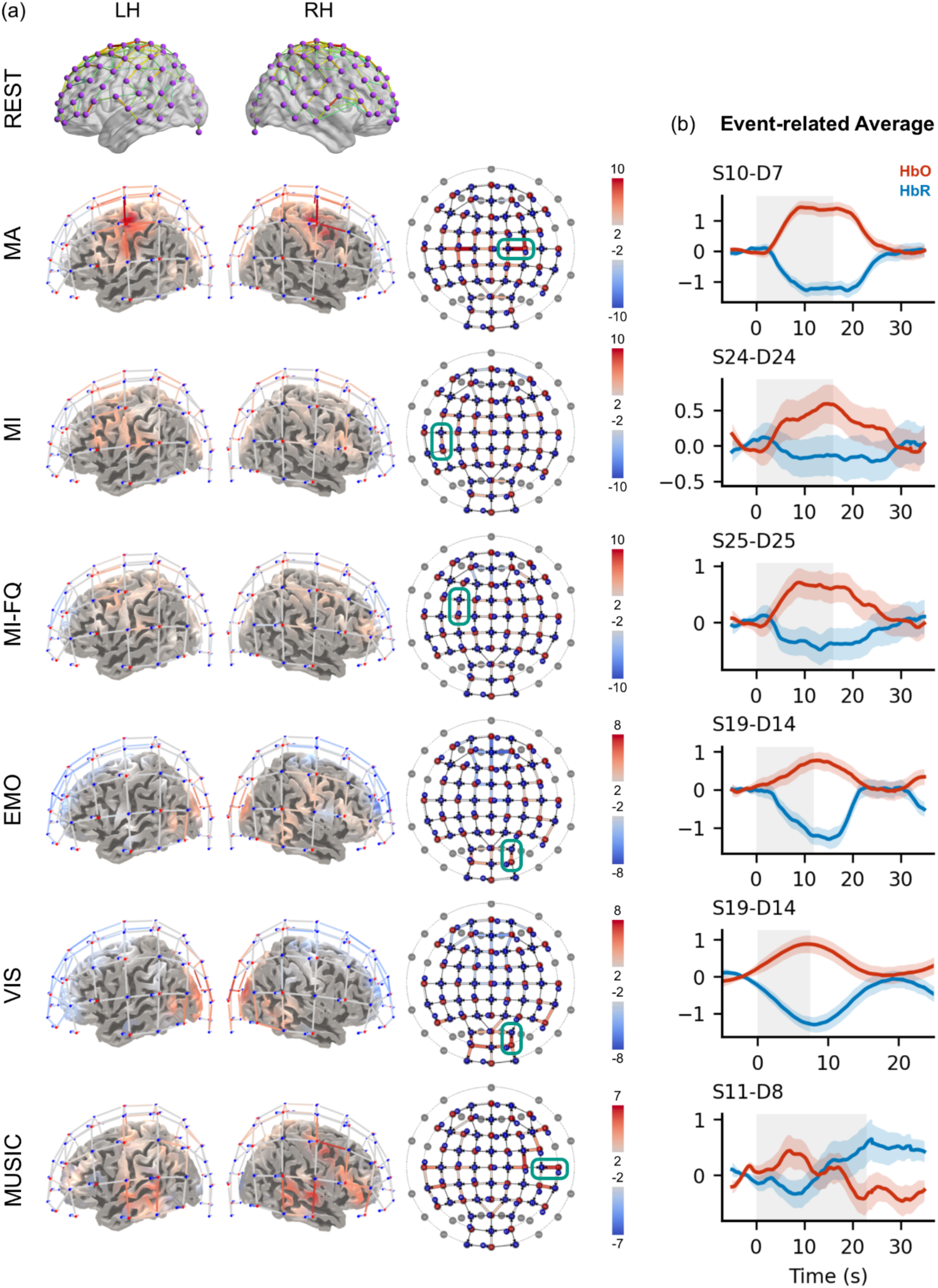
(a) Connectivity map of the resting state task (top 5% strongest HbO connections at the group level) and RFX GLM results (t-maps) of the HbO data for each active task (n = 22), thresholded at *p* ≤ .05. Results are displayed as exploratory surface projections on the LH = left hemisphere, RH = right hemisphere, and flat maps. For the visual, music and emotion task, all conditions are jointly contrasted with the baseline period. Channels with the highest t-value are highlighted on the flat maps in green for each task and the respective Event-related Averages of HbO and HbR are shown in (b). Grey shading indicates the task period. For the emotion task, the “Neutral-Happy” response is displayed. For the Visual Task, the “Threatening” category was chosen as an example. The music task shows the response to a “High-Valence, High-Arousal” stimulus. REST = Resting State, MA = Motor Action, MI = Motor Imagery, MI-FQ = Motor Imagery with frequency change, EMO = Emotion, VIS = Visual.

For the motor action task, exploratory surface-projected RFX GLM analyses of the HbO data (*p* ≤ .05, uncorrected) showed activity in the left and right pre-motor and supplementary motor areas, as well as in the bilateral primary motor and sensory cortices for the task > baseline contrast. The motor imagery task > baseline contrast (*p* ≤ .05, uncorrected) revealed activity in the left pre-motor and supplementary motor areas, with activation extending laterally into the left supramarginal gyrus, left frontal eye fields, and left and right Broca’s area, and weak deactivation in frontal areas. For the motor imagery task including frequency changes in the imagined movement, the task > baseline contrast (*p* ≤ .05, uncorrected) showed weaker amplitudes than the other two tasks and revealed activation in bilateral pre-motor and supplementary motor areas, as well as the right anterior frontal cortex, and deactivation in anterior prefrontal areas.

GLM analysis (*p* ≤ .05, uncorrected) of the emotion task revealed condition-related differences in cortical activation during emotional face processing. For the sad > neutral contrast, activation increases were observed in a distributed frontal-posterior temporal network. In the left hemisphere, effects involved inferior frontal and frontal opercular cortex, frontocentral cortex, and posterior temporal cortex centered on the posterior superior temporal sulcus (pSTS), extending toward adjacent inferior parietal regions. Additional effects were present in the right hemisphere over the frontocentral and posterior temporoparietal cortex. Thus, the sad > neutral contrast was characterized by relatively widespread bilateral activity, with a left-hemispheric predominance. For the happy > neutral contrast, activation was more restricted and more left-lateralized. The main effect was located in the left posterior temporal cortex, again centered on the pSTS region, with limited extension toward adjacent posterior temporoparietal/inferior parietal cortex. Smaller effects were also observed in the left lateral frontal cortex. No significant effects were found for the happy > sad contrast.

For the visual task, the task > baseline contrast (*p* ≤ .05, uncorrected) showed activity in the bilateral secondary visual and visual association cortices. This activity was accompanied by widespread deactivation, especially in dorsolateral prefrontal regions.

The music task > baseline contrast (*p* ≤ .05, uncorrected) showed activation in the bilateral temporal auditory cortices, superior temporal gyri, right pre-motor and supplementary motor areas and right homologue of Broca’s area. Surface-projected connectivity and t-maps for the HbR data can be found in Appendix Fig. S3 for all tasks.

### 3.3 Signal quality of physiological data and task effects on physiology

Runs in which physiological sensors showed clear connection issues as determined by visual inspection were excluded from the analyses. Excluded runs for the same sensor type often occurred in the same subject, with 46.67% of respiratory, 82.61% of HR, 73.08% of GSR, 50% of ECG, and 28.57% of sensor issues occurring in subjects who had sensor issues in more than one run. These results suggest that sensors detached during measurement, resulting in consistently bad/missing data for the rest of the run. PPG issues were never found in the same run. Figure 7 (a) shows the signal quality and task effects of the physiological measures. Physiological data showed overall satisfactory data quality with SNR of the physio data of mean_SNR-Resp_= 43.39 ± 4.2 dB; mean_SNR-ECG_= 20.26 ± 12.59 dB; mean_SNR-PPG_= 44.77 ± 7.81 dB; mean_SNR-GSR_= 25.69 ± 9.16 dB, and with average heart rate measured across all participants and runs of mean_HR_ = 72.81 ± 3.14 bpm, and the breathing rate of mean_BR_= 14.89 ± 3.69 rpm. Figure 7 (b) suggests that mean heart rate was relatively stable across conditions, although there was noticeable inter-individual variability. Breathing rate appeared to be lowest for the resting state and slightly higher during task conditions, with some variation across tasks and participants. No statistical tests were conducted to formally assess condition differences.

**Fig. 7.**
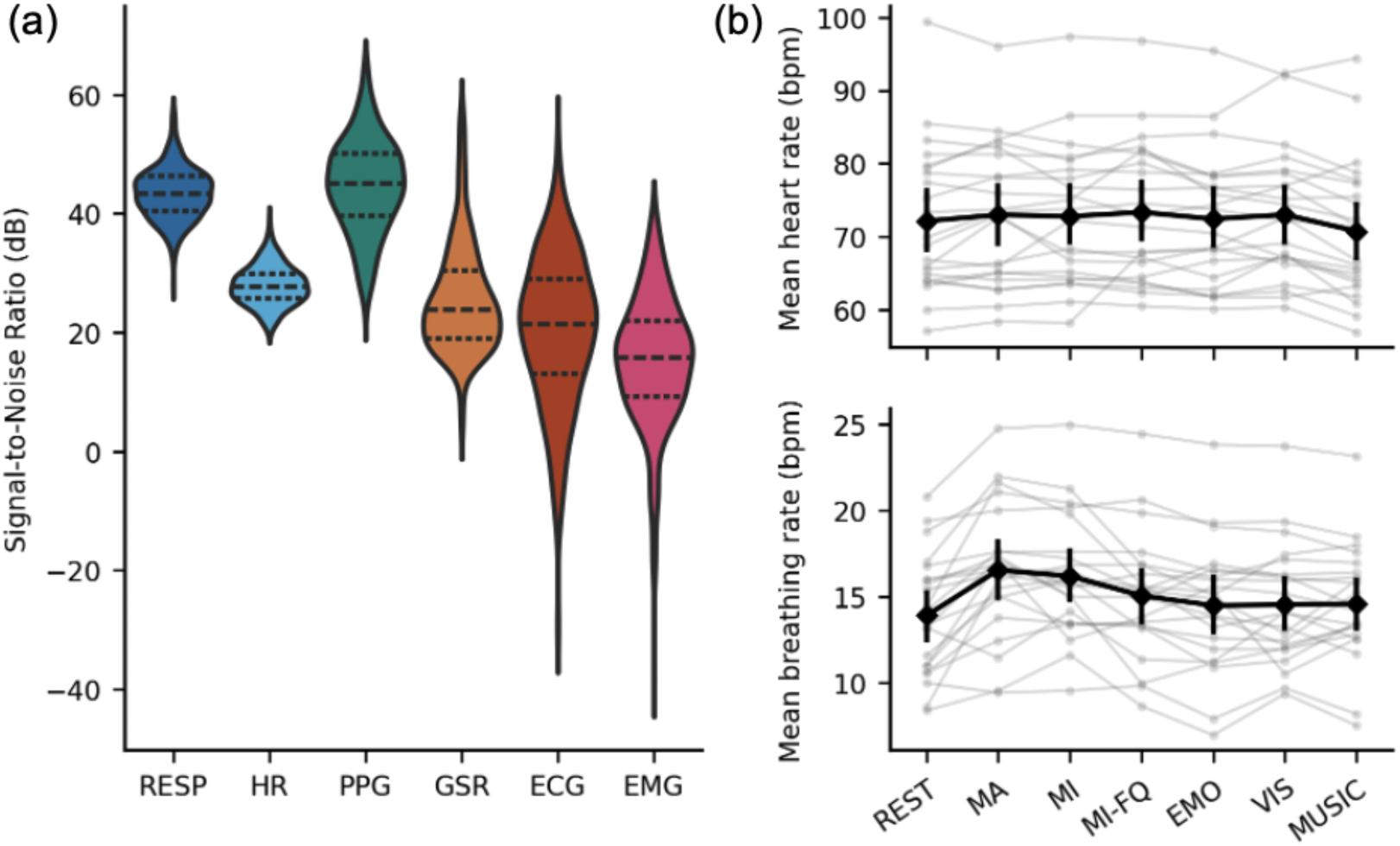
(a) Physiological data quality across all participants (n = 57) and runs, excluding runs with sensor issues. Dashed and dotted lines represent quartiles. (b) Mean heart (beats per minute) and breathing rate (breaths per minute) across tasks (n = 22). Grey lines show the means per subject. Black lines indicate group-level means, with error bars representing the 95% confidence interval.

## 4 Discussion

fNIRS is increasingly used to study brain activity in naturalistic and personalized settings, but task-evoked responses often show substantial inter-individual variability. This variability may reflect meaningful neurovascular differences, but it may also arise from signal quality differences and systemic physiological confounds. In this study, we therefore examined whether channel-level signal quality contributes to task-related fNIRS activation estimates and present a large, openly available dataset designed to support systematic evaluation of signal quality and physiology-correction methods.

### 4.1 SNR, CV, SCI and cSNR capture distinct features of fNIRS signal quality

We evaluated signal quality with four different measures: SNR, CV, SCI, and cSNR. As expected for SNR and CV due to their inverse mathematical relationship, these values were highly negatively correlated. The other quality metrics showed only low to medium correlations with each other. These findings show that each measure captures distinct signal properties. Reporting several quality metrics may thus give a more comprehensive overview of signal quality in a dataset. Further, signal quality was systematically related to channel length, although the direction and strength of the association differed across metrics. The strongest relationship was observed for SNR, which decreased substantially with increasing source-detector distance. Increased channel length was also associated with lower SCI and higher CV. Results for these three quality metrics are in line with expectations, as increased source-detector separation is generally associated with reduced light intensity and lower signal-to-noise ratio.^4,29,55^ Unexpectedly, channel length was positively associated with cSNR, such that shorter channels exhibited lower cSNR values. Because cSNR reflects the ratio of cardiac-band power to noise-band power, lower cSNR values indicate weaker cardiac coupling relative to noise. This pattern is counterintuitive, as shorter channels, particularly short-separation channels, are typically expected to be more sensitive to superficial physiological signals, including cardiac activity. Accordingly, these channels would be expected to show stronger cardiac contributions, and therefore higher rather than lower cSNR values. Possibly, this is related to the measure capturing relative dominance of cardiac-band power over flanking noise bands, which would lead to higher cSNR values if shorter channels contain elevated non-cardiac (physiological) noise compared to regular channels, thereby lowering cSNR.

The cSNR metric implemented here is conceptually similar to cardiac pulse SNR measures reported in high-density applications of fNIRS, in which raw optical signal quality is quantified as cardiac- or pulse-band power relative to flanking/background spectral power.^56–58^ While these approaches commonly use a broader pulse band and bandwidth-scaled flanking noise estimates across the cap, the present implementation used fixed cardiac and noise bands and computed a channel-wise dB ratio averaged across wavelengths. Although beyond the scope of the present study, future work should directly compare the current cSNR approach with cardiac pulse SNR metrics used in high-density fNIRS literature. Such analyses could clarify whether the unexpected positive association between channel length and cSNR reported here reflects a property of the present metric specifically, or whether a similar relationship is also observed when signal quality is quantified using established pulse SNR measures.

### 4.2 Signal quality partially explains task-evoked fNIRS responses

To assess whether signal quality differences might contribute to large inter-subject variability in fNIRS response previously reported in the literature,^14–17^ we investigated the effects of signal quality on task-evoked hemodynamic changes in a motor action task. The motor action task was chosen because this task showed reliable, clearly localized responses in the fNIRS signal. We found significant association of all signal quality metrics to task-related fNIRS responses, suggesting that signal quality is not merely a technical nuisance addressed through channel rejection, but may also influence the magnitude of task-related effects in the retained data. For the wavelength-specific measures, the direction of the associations was largely compatible with better signal quality being associated with more clearly expressed task-related responses: higher SNR and lower CV were generally associated with more positive HbO and/or more negative HbR estimates. A similar pattern was evident for SCI in HbO and for cSNR in several channels, particularly S25-D24, where higher cSNR was associated with both a more positive HbO estimate and a more negative HbR estimate. In contrast, SCI associations with HbT were directionally mixed, and cSNR showed no significant HbT associations. The recurring involvement of several channels across quality metrics suggests that measurement quality may have attenuated or obscured task-related responses in portions of the sensorimotor montage. While we only investigated signal quality effects on a motor action paradigm here, we expect similar effects in other tasks. Notably, these effects were found after SCI-based channel rejection, suggesting that signal quality influences on task activations persist even after quality control procedures. Since this was not investigated in the current study, it remains unclear whether an analogous relationship would be observed after CV-based channel rejection. As evident from the correlation between the two metrics, SCI- and CV-based criteria capture partly distinct aspects of data quality, with SCI indexing cardiac-band coupling across wavelengths and CV reflecting relative signal variability. Therefore, CV-based rejection may attenuate quality-related covariance with task-evoked activation if the effect is driven by unstable or noisy channels. Future work might compare SCI-, CV- or other rejection strategies within the same dataset to determine whether quality-related covariance with task-evoked activation persists across preprocessing choices. Such analyses could provide further insight into how signal-quality differences contribute to inter-subject variability and inform preprocessing strategies for reducing quality-related variance that may obscure task-evoked effects.

Although the present analysis cannot determine whether signal quality causally altered the measured response, the observed association indicates that signal quality may (partially) explain inter-subject variability in task-related activation estimates. Such effects are particularly important in the development of personalized fNIRS applications, as well as studies comparing conditions, cortical regions, or participant groups, where systematic differences in optode-scalp coupling or signal robustness could confound apparent activation differences. Thus, signal quality should not be treated solely as a binary exclusion criterion, but also as a potential source of variance in the retained data.

### 4.3 Functional validation of the MULPA dataset

Exploratory analyses of task effects in the fNIRS signal showed distinct cortical response profiles for the different tasks. In all motor tasks, activity was localized in the pre-motor, supplementary, and primary motor areas. Although the tasks are referred to here as motor tasks, they were sensorimotor in nature. Execution and imagery of movement can involve not only motor planning and primary motor regions, but also somatosensory processing related to proprioceptive, tactile, and body-state representations.^59^ Accordingly, activation was observed not only in expected motor areas, but also in primary sensory cortices. Mental imagery motor tasks elicited weaker activations than the motor action task, with increasing task complexity further reducing activation in motor cortices but resulting in de-activations in dorsolateral prefrontal areas. These findings are consistent with previous fNIRS studies demonstrating motor-related hemodynamic responses during both executed and imagined movements and with broader neuroimaging evidence identifying premotor and supplementary motor areas as core components active in motor-imagery paradigms.^60–62^ In the emotion task, sad faces elicited a broader response involving left inferior frontal/frontal opercular, frontocentral, and posterior temporal regions, with additional right frontocentral and posterior temporoparietal involvement, in particular the pSTS. These findings are consistent with facial-emotion recognition relying on a distributed cortical system rather than a single specialized region,^63^ and with evidence implicating the pSTS and inferior frontal operculum in emotion discrimination.^64,65^ In the visual task, activation was mainly observed in occipital regions, consistent with previous fNIRS studies demonstrating increased visual-cortical responses during photic or pattern-based stimulation.^66–68^ In addition, we observed reduced responses in frontoparietal regions during visual stimulation. This pattern may be explained by task-induced suppression of internally oriented processing during externally directed visual engagement, as reported in functional neuroimaging studies of visual tasks.^69,70^ The music task elicited activation in bilateral temporal auditory regions, including channels localized to the superior temporal gyri, consistent with the established role of auditory association cortices in the processing of complex musical sounds and with previous fNIRS studies demonstrating temporal cortical responses during music listening.^71–73^ Given the limited depth sensitivity of fNIRS, the temporal findings are most cautiously interpreted as reflecting auditory cortical and superior temporal involvement rather than definitive localization to primary auditory cortex.

In summary, all task-induced activation changes were observed in expected areas for each task, supporting the suitability of this dataset for investigating further empirical research questions related to the task battery. However, as these effects were observed at an uncorrected threshold, the findings should be interpreted as preliminary.

### 4.4 Signal quality of physiological data and task effects on physiology

In addition to task-based fNIRS recordings, the present dataset includes a comprehensive set of peripheral physiological measures. To characterize the quality of these recordings, we report the SNR across participants and runs. We also provide descriptive summaries of heart rate and respiratory rate across tasks. These analyses are intended to document the availability and validity of the physiological recordings rather than to test task-related physiological effects. Accordingly, no inferential statistical analyses were performed, and no conclusions regarding task-induced differences in peripheral physiology can be drawn from this study. However, the availability of these comprehensive physiological recordings makes the dataset a valuable resource for future work on how systemic physiology influences fNIRS signals and for the development and evaluation of physiology-informed correction methods.

Although the presented, openly available dataset is relatively large by fNIRS standards, its scale remains modest compared with resources available for other neuroimaging modalities, such as large functional magnetic resonance imaging (fMRI) initiatives including the Human Connectome Project.^75^ Moreover, the study was not preregistered, and the sampling plan was not determined a priori based on formal power calculations, which limits the strength of confirmatory inferences. However, to our knowledge, this is the first comprehensive investigation with regards to physiological effects with shared data and code, allowing for full reproducibility, systematic data exploration and exploiting analytical degrees of freedom to determine optimal processing and analysis steps.^76^ Future work would benefit from coordinated multi-site efforts to establish large, standardized fNIRS datasets that include short-separation channels, concurrent physiological recording, and a large cognitive task battery. Such datasets would provide an important resource for benchmarking preprocessing pipelines, developing and validating physiological noise-correction methods, and enabling more data-intensive approaches, including machine learning models for quality assessment, artifact correction, and task decoding.

## 5 Conclusion

In conclusion, the present study demonstrates that fNIRS quality metrics capture distinct, complementary characteristics of the signal and are systematically associated with task-related activation estimates. These findings support the view that signal quality should be reported comprehensively and may partially explain inter-individual differences in task-evoked activation patterns. It may thus be advisable to not only consider data quality during channel rejection, but also as a potential source of variance in statistical analyses. Further, the presented dataset includes extensive short-channel and peripheral physiological recordings, making it a valuable resource for evaluating signal quality effects and developing physiology correction approaches. By openly providing this multimodal dataset, the study aims to support more transparent, reproducible, and physiologically informed fNIRS research, particularly for applications in which individual-level activation estimates are central.

## 6 Code and Data Availability

### Disclosures

AC, ML, and RG are employees of Brain Innovation B.V. (Maastricht, The Netherlands). This has not affected the content or quality of this work. All other authors declare no conflict of interest.

### Code, Data, and Materials

All data, as well as task presentation scripts and preprocessing pipelines presented in this manuscript are openly available in “The MULtiple PAradigm (MULPA) Dataset” repository at https://zenodo.org/records/21033499 under the Creative Commons Attribution-NonCommercial 4.0 International License (CC BY-NC 4.0).

## Acknowledgments

The authors acknowledge funding through The Netherlands Organization for Scientific Research (NWO; Vidi-Grant No. VI.Vidi.191.210 to B.S.). D.M.A.M is supported by a Junior Principal Investigator (JPI) fellowship of RWTH Aachen university funded by the Excellence Strategy of the Federal Government and the Laender (Grant No. JPI074-21). S.R. is supported by the Cusanuswerk scholarship foundation. J.P., B.D., and T.S. were supported by the Portuguese Foundation for Science and Technology (FCT; Grants No. 2020.04899.BD, CEECINST/00117/2021/CP2784/CT0002, and CEECINSTLA/00026/2022/CP2919/CT0001). GPT-5.5 Thinking was used to assist with data visualization code, to provide feedback on paragraph structure and to edit grammar for portions of the manuscript, as well as to create appendix figures visualizing the calculation of cSNR and Euclidean distance. Nano Banana Pro was used to generate the illustration of the human wearing fNIRS and physiology sensors in Fig 1. All content was reviewed, verified, and approved by the authors, who take full responsibility for the manuscript.

**Sophie Raible** is a PhD candidate at the Department of Cognitive Neuroscience of Maastricht University. She received her BSc and MSc degrees in Psychology and Neuroscience with distinction from Maastricht University in 2019 and 2022, respectively. Her research focuses on (concurrent) fNIRS and fMRI, visual perception, mental imagery and brain-computer interfaces.

**João Pereira** is a Biomedical Engineer and a PhD candidate at the University of Coimbra and Maastricht University, with current work being developed at Coimbra Institute for Biomedical Imaging and Translational Research (CIBIT). His research focuses on the development and validation of neurorehabilitative strategies based on multimodal imaging (fMRI and fNIRS).

**David M.A. Mehler** is a physician-scientist and heads the Applied Computational Neuroscience lab at Uniklinik RWTH Aachen, Germany. The research of his group focuses on non-invasive brain-computer-interfaces (neurofeedback), metaresearch for neurotechnology, as well as machine-learning based big data approaches to investigate clinical and biological markers and predictors for psychiatric and neurological diseases.

**Simon H. Kohl** is a postdoctoral researcher at Forschungszentrum Jülich and RWTH Aachen University Hospital, Germany. His work focuses on neurofeedback, as well as the development and evidence-based implementation of digital health solutions to improve mental health care for children and adolescents.

**Miguel Castelo-Branco** is a full professor at the University of Coimbra and Director of the Coimbra Institute for Biomedical Imaging and Translational Research. His main research interests are visual, cognitive and clinical neuroscience, and in therapeutic pharmacological and non pharmacological approaches such as neurofeedback.

**Michael Lührs** is a postdoctoral researcher in the Department of Cognitive Neuroscience at Maastricht University and Brain Innovation B.V. His main research focus is the advancement of neurofeedback tools and other real-time applications using fMRI and fNIRS.

**Bettina Sorger** is an associate professor in the Department of Cognitive Neuroscience at Maastricht University, The Netherlands. Her main research interests include motor-independent communication/environmental control and neurofeedback (therapy) - both based on brain hemodynamics as measured with fNIRS and fMRI.

Biographies for the other authors are not available.

## References

1. F. F. Jöbsis, “Noninvasive, infrared monitoring of cerebral and myocardial oxygen sufficiency and circulatory parameters,” Science 198(4323), 1264–1267, New York, N.Y. (1977) [doi:10.1126/science.929199].

2. M. Ferrari and V. Quaresima, “A brief review on the history of human functional near-infrared spectroscopy (fNIRS) development and fields of application,” NeuroImage 63(2), 921–935 (2012) [doi:10.1016/j.neuroimage.2012.03.049].

3. M. A. Yücel et al., “Functional Near Infrared Spectroscopy: Enabling Routine Functional Brain Imaging,” Curr. Opin. Biomed. Eng. 4, 78–86 (2017) [doi:10.1016/j.cobme.2017.09.011].

4. P. Pinti et al., “The present and future use of functional near-infrared spectroscopy (fNIRS) for cognitive neuroscience,” Ann. N. Y. Acad. Sci. 1464(1), 5–29 (2020) [doi:10.1111/nyas.13948].

5. D. A. Boas et al., “Twenty years of functional near-infrared spectroscopy: introduction for the special issue,” NeuroImage 85 **Pt** **1**, 1–5 (2014) [doi:10.1016/j.neuroimage.2013.11.033].

6. Y. Gong et al., “The Use of Functional Near-Infrared Spectroscopy (fNIRS) for Monitoring Brain Function, Predicting Outcomes, and Evaluating Rehabilitative Interventional Responses in Poststroke Patients With Upper Limb Hemiplegia: A Systematic Review,” IEEE J. Sel. Top. Quantum Electron. 31(4: Adv. in Neurophoton. for Non-Inv. Brain Mon.), 1–10 (2025) [doi:10.1109/JSTQE.2025.3563153].

7. M. Mihara and I. Miyai, “Review of functional near-infrared spectroscopy in neurorehabilitation,” Neurophotonics 3(3), 031414 (2016) [doi:10.1117/1.NPh.3.3.031414].

8. C. Huo et al., “A review on functional near-infrared spectroscopy and application in stroke rehabilitation,” Med. Nov. Technol. Devices 11, 100064 (2021) [doi:10.1016/j.medntd.2021.100064].

9. N. Naseer and K.-S. Hong, “fNIRS-based brain-computer interfaces: a review,” Front. Hum. Neurosci. 9, 3 (2015) [doi:10.3389/fnhum.2015.00003].

10. K. Paulmurugan et al., “Brain–Computer Interfacing Using Functional Near-Infrared Spectroscopy (fNIRS),” Biosensors 11(10), 389 (2021) [doi:10.3390/bios11100389].

11. S. H. Kohl et al., “The Potential of Functional Near-Infrared Spectroscopy-Based Neurofeedback—A Systematic Review and Recommendations for Best Practice,” Front. Neurosci. 14, Frontiers (2020) [doi:10.3389/fnins.2020.00594].

12. F. Klein et al., “From lab to life: challenges and perspectives of fNIRS for hemodynamic-based neurofeedback in real-world environments,” Philos. Trans. R. Soc. B Biol. Sci. 379(1915), 20230087 (2024) [doi:10.1098/rstb.2023.0087].

13. S. R. Soekadar et al., “Optical brain imaging and its application to neurofeedback,” NeuroImage Clin. 30, 102577 (2021) [doi:10.1016/j.nicl.2021.102577].

14. H. Zohdi, F. Scholkmann, and U. Wolf, “Individual Differences in Hemodynamic Responses Measured on the Head Due to a Long-Term Stimulation Involving Colored Light Exposure and a Cognitive Task: A SPA-fNIRS Study,” Brain Sci. 11(1), 54 (2021) [doi:10.3390/brainsci11010054].

15. H. Zohdi, F. Scholkmann, and U. Wolf, “Long-term blue light exposure changes frontal and occipital cerebral hemodynamics: Not all subjects react the same,” in Oxygen transport to tissue XLII, E. M. Nemoto et al., Eds., pp. 217–222, Springer International Publishing, Cham (2021) [doi:10.1007/978-3-030-48238-1_34].

16. L. Minati et al., “Variability comparison of simultaneous brain near-infrared spectroscopy (NIRS) and functional MRI (fMRI) during visual stimulation,” J. Med. Eng. Technol. 35(0), 370–376 (2011) [doi:10.3109/03091902.2011.595533].

17. D. G. Wyser et al., “Characterizing reproducibility of cerebral hemodynamic responses when applying short-channel regression in functional near-infrared spectroscopy,” Neurophotonics 9(01) (2022) [doi:10.1117/1.NPh.9.1.015004].

18. I. Tachtsidis and F. Scholkmann, “False positives and false negatives in functional near-infrared spectroscopy: issues, challenges, and the way forward,” Neurophotonics 3(3), 031405 (2016) [doi:10.1117/1.NPh.3.3.031405].

19. A. Abdalmalak et al., “Effects of Systemic Physiology on Mapping Resting-State Networks Using Functional Near-Infrared Spectroscopy,” Front. Neurosci. 16, Frontiers (2022) [doi:10.3389/fnins.2022.803297].

20. F. Scholkmann et al., “Systemic physiology augmented functional near-infrared spectroscopy: a powerful approach to study the embodied human brain,” Neurophotonics 9(3), 030801 (2022) [doi:10.1117/1.NPh.9.3.030801].

21. B. Wang, A. M. Goodpaster, and M. A. Kennedy, “Coefficient of variation, signal-to-noise ratio, and effects of normalization in validation of biomarkers from NMR-based metabonomics studies,” Chemom. Intell. Lab. Syst. 128, 9–16 (2013) [doi:10.1016/j.chemolab.2013.07.007].

22. M. A. Yücel et al., “Best practices for fNIRS publications,” Neurophotonics 8(1), 012101 (2021) [doi:10.1117/1.NPh.8.1.012101].

23. L. Pollonini et al., “Auditory cortex activation to natural speech and simulated cochlear implant speech measured with functional near-infrared spectroscopy,” Hear. Res. 309, 84–93 (2014) [doi:10.1016/j.heares.2013.11.007].

24. L. Pollonini, H. Bortfeld, and J. S. Oghalai, “PHOEBE: a method for real time mapping of optodes-scalp coupling in functional near-infrared spectroscopy,” Biomed. Opt. Express 7(12), 5104–5119 (2016) [doi:10.1364/BOE.7.005104].

25. K. Tripathy et al., “Decoding visual information from high-density diffuse optical tomography neuroimaging data,” NeuroImage 226, 117516 (2021) [doi:10.1016/j.neuroimage.2020.117516].

26. M. S. Sappia et al., “Signal quality index: an algorithm for quantitative assessment of functional near infrared spectroscopy signal quality,” Biomed. Opt. Express 11(11), 6732–6754 (2020) [doi:10.1364/BOE.409317].

27. F. Scholkmann et al., “A review on continuous wave functional near-infrared spectroscopy and imaging instrumentation and methodology,” NeuroImage 85, 6–27 (2014) [doi:10.1016/j.neuroimage.2013.05.004].

28. M. A. Yücel et al., “Short separation regression improves statistical significance and better localizes the hemodynamic response obtained by near-infrared spectroscopy for tasks with differing autonomic responses,” Neurophotonics 2(3), 035005 (2015) [doi:10.1117/1.NPh.2.3.035005].

29. S. Brigadoi and R. J. Cooper, “How short is short? Optimum source–detector distance for short-separation channels in functional near-infrared spectroscopy,” Neurophotonics 2(2), 025005 (2015) [doi:10.1117/1.NPh.2.2.025005].

30. L. Gagnon et al., “Short separation channel location impacts the performance of short channel regression in NIRS,” NeuroImage 59(3), 2518–2528 (2012) [doi:10.1016/j.neuroimage.2011.08.095].

31. L. Gagnon et al., “Improved recovery of the hemodynamic response in diffuse optical imaging using short optode separations and state-space modeling,” NeuroImage 56(3), 1362–1371 (2011) [doi:10.1016/j.neuroimage.2011.03.001].

32. A. von Lühmann et al., “Improved physiological noise regression in fNIRS: A multimodal extension of the General Linear Model using temporally embedded Canonical Correlation Analysis,” NeuroImage 208, 116472 (2020) [doi:10.1016/j.neuroimage.2019.116472].

33. H. Santosa et al., “Quantitative comparison of correction techniques for removing systemic physiological signal in functional near-infrared spectroscopy studies,” Neurophotonics 7(3), 035009 (2020) [doi:10.1117/1.NPh.7.3.035009].

34. Q. Zhang, E. N. Brown, and G. E. Strangman, “Adaptive filtering for global interference cancellation and real-time recovery of evoked brain activity: a Monte Carlo simulation study,” J. Biomed. Opt. 12(4), 044014, SPIE (2007) [doi:10.1117/1.2754714].

35. T. Yamada, S. Umeyama, and K. Matsuda, “Multidistance probe arrangement to eliminate artifacts in functional near-infrared spectroscopy,” J. Biomed. Opt. 14(6), 064034, SPIE (2009) [doi:10.1117/1.3275469].

36. S. Umeyama and T. Yamada, “Monte Carlo study of global interference cancellation by multidistance measurement of near-infrared spectroscopy,” J. Biomed. Opt. 14(6), 064025 (2009) [doi:10.1117/1.3275466].

37. D. Lakens, “Sample Size Justification,” Collabra Psychol. 8(1), D. van Ravenzwaaij, Ed., 33267 (2022) [doi:10.1525/collabra.33267].

38. G. A. Zimeo Morais, J. B. Balardin, and J. R. Sato, “fNIRS Optodes’ Location Decider (fOLD): a toolbox for probe arrangement guided by brain regions-of-interest,” 1, Sci. Rep. 8(1), 3341, Nature Publishing Group (2018) [doi:10.1038/s41598-018-21716-z].

39. S. E. Williams et al., “Further validation and development of the movement imagery questionnaire,” J. Sport Exerc. Psychol. 34(5), 621–646 (2012) [doi:10.1123/jsep.34.5.621].

40. D. M. McNair, M. Lorr, and L. F. Droppleman, *Profile of mood states*, Educational and Industrial Testing Service, San Diego, Calif. (1971).

41. X. Wu et al., “Acquisition time for functional near-infrared spectroscopy resting-state functional connectivity in assessing autism,” Neurophotonics 9(4), 045007 (2022) [doi:10.1117/1.NPh.9.4.045007].

42. S. Geng et al., “Effect of Resting-State fNIRS Scanning Duration on Functional Brain Connectivity and Graph Theory Metrics of Brain Network,” Front. Neurosci. 11, 392 (2017) [doi:10.3389/fnins.2017.00392].

43. J. Pereira et al., “Self-Modulation of Premotor Cortex Interhemispheric Connectivity in a Real-Time Functional Magnetic Resonance Imaging Neurofeedback Study Using an Adaptive Approach,” Brain Connect. 9(9), 662–672 (2019) [doi:10.1089/brain.2019.0697].

44. P. Lucey et al., “The Extended Cohn-Kanade Dataset (CK+): A complete dataset for action unit and emotion-specified expression,” in 2010 IEEE Computer Society Conference on Computer Vision and Pattern Recognition - Workshops, pp. 94–101, IEEE, San Francisco, CA, USA (2010) [doi:10.1109/CVPRW.2010.5543262].

45. I. Almeida, M. Van Asselen, and M. Castelo-Branco, “The role of the amygdala and the basal ganglia in visual processing of central vs. peripheral emotional content,” Neuropsychologia 51(11), 2120–2129 (2013) [doi:10.1016/j.neuropsychologia.2013.07.007].

46. R. Panda, R. Malheiro, and R. P. Paiva, “Novel Audio Features for Music Emotion Recognition,” IEEE Trans. Affect. Comput. 11(4), 614–626 (2020) [doi:10.1109/TAFFC.2018.2820691].

47. J. A. Russell, “A circumplex model of affect,” J. Pers. Soc. Psychol. 39(6), 1161–1178, American Psychological Association, US (1980) [doi:10.1037/h0077714].

48. F. A. Fishburn et al., “Temporal Derivative Distribution Repair (TDDR): A motion correction method for fNIRS,” NeuroImage 184, 171–179 (2019) [doi:10.1016/j.neuroimage.2018.09.025].

49. Y.-J. Zhang, “Detecting resting-state functional connectivity in the language system using functional near-infrared spectroscopy,” J. Biomed. Opt. 15(4), 047003 (2010) [doi:10.1117/1.3462973].

50. C.-M. Lu et al., “Use of fNIRS to assess resting state functional connectivity,” J. Neurosci. Methods 186(2), 242–249 (2010) [doi:10.1016/j.jneumeth.2009.11.010].

51. L. Duan, Y.-J. Zhang, and C.-Z. Zhu, “Quantitative comparison of resting-state functional connectivity derived from fNIRS and fMRI: a simultaneous recording study,” NeuroImage 60(4), 2008–2018 (2012) [doi:10.1016/j.neuroimage.2012.02.014].

52. K. Smitha et al., “Resting state fMRI: A review on methods in resting state connectivity analysis and resting state networks,” Neuroradiol. J. 30(4), 305–317, SAGE Publications Ltd (2017) [doi:10.1177/1971400917697342].

53. H. Zhang et al., “Functional connectivity as revealed by independent component analysis of resting-state fNIRS measurements,” NeuroImage 51(3), 1150–1161 (2010) [doi:10.1016/j.neuroimage.2010.02.080].

54. C. Notte, C. Alionte, and C. D. Strubakos, “The efficacy and methodology of using near-infrared spectroscopy to determine resting-state brain networks,” J. Neurophysiol. 131(4), 668–677 (2024) [doi:10.1152/jn.00357.2023].

55. G. E. Strangman, Z. Li, and Q. Zhang, “Depth Sensitivity and Source-Detector Separations for Near Infrared Spectroscopy Based on the Colin27 Brain Template,” PLoS ONE 8(8), X.-N. Zuo, Ed., e66319 (2013) [doi:10.1371/journal.pone.0066319].

56. M. Fogarty et al., “Functional brain mapping using whole-head very high-density diffuse optical tomography,” Imaging Neurosci. 3, IMAG.a.54 (2025) [doi:10.1162/IMAG.a.54].

57. A. Sherafati et al., “Prefrontal cortex supports speech perception in listeners with cochlear implants,” eLife 11, T. D. Griffiths et al., Eds., e75323, eLife Sciences Publications, Ltd (2022) [doi:10.7554/eLife.75323].

58. K. Tripathy et al., “Mapping brain function in adults and young children during naturalistic viewing with high-density diffuse optical tomography,” Hum. Brain Mapp. 45(7), e26684 (2024) [doi:10.1002/hbm.26684].

59. R. M. Hardwick et al., “Neural correlates of action: Comparing meta-analyses of imagery, observation, and execution,” Neurosci. Biobehav. Rev. 94, 31–44 (2018) [doi:10.1016/j.neubiorev.2018.08.003].

60. F. Klein et al., “fMRI-based validation of continuous-wave fNIRS of supplementary motor area activation during motor execution and motor imagery,” Sci. Rep. 12(1), 3570, Nature Publishing Group (2022) [doi:10.1038/s41598-022-06519-7].

61. J. Pereira et al., “Multimodal assessment of the spatial correspondence between fNIRS and fMRI hemodynamic responses in motor tasks,” Sci. Rep. 13 (2023) [doi:10.1038/s41598-023-29123-9].

62. R. M. Hardwick et al., “Neural correlates of action: Comparing meta-analyses of imagery, observation, and execution,” Neurosci. Biobehav. Rev. 94, 31–44 (2018) [doi:10.1016/j.neubiorev.2018.08.003].

63. R. Adolphs et al., “Cortical systems for the recognition of emotion in facial expressions,” J. Neurosci. Off. J. Soc. Neurosci. 16(23), 7678–7687 (1996) [doi:10.1523/JNEUROSCI.16-23-07678.1996].

64. B. Direito et al., “Targeting dynamic facial processing mechanisms in superior temporal sulcus using a novel fMRI neurofeedback target,” Neuroscience 406, 97–108 (2019) [doi:10.1016/j.neuroscience.2019.02.024].

65. I. Iarrobino et al., “Right and left inferior frontal opercula are involved in discriminating angry and sad facial expressions,” Brain Stimulat. 14(3), 607–615 (2021) [doi:10.1016/j.brs.2021.03.014].

66. T. Kato et al., “Human Visual Cortical Function during Photic Stimulation Monitoring by Means of near-Infrared Spectroscopy,” J. Cereb. Blood Flow Metab. 13(3), 516–520, SAGE Publications Ltd STM (1993) [doi:10.1038/jcbfm.1993.66].

67. M. M. Plichta et al., “Event-Related Visual versus Blocked Motor Task: Detection of Specific Cortical Activation Patterns with Functional Near-Infrared Spectroscopy,” Neuropsychobiology 53(2), 77–82 (2006) [doi:10.1159/000091723].

68. A. T. Eggebrecht et al., “A quantitative spatial comparison of high-density diffuse optical tomography and fMRI cortical mapping,” NeuroImage 61(4), 1120–1128 (2012) [doi:10.1016/j.neuroimage.2012.01.124].

69. K. D. Singh and I. P. Fawcett, “Transient and linearly graded deactivation of the human default-mode network by a visual detection task,” NeuroImage 41(1), 100–112 (2008) [doi:10.1016/j.neuroimage.2008.01.051].

70. M. E. Raichle et al., “A default mode of brain function,” Proc. Natl. Acad. Sci. 98(2), 676–682, Proceedings of the National Academy of Sciences (2001) [doi:10.1073/pnas.98.2.676].

71. M. M. Plichta et al., “Auditory cortex activation is modulated by emotion: A functional near-infrared spectroscopy (fNIRS) study,” NeuroImage 55(3), 1200–1207 (2011) [doi:10.1016/j.neuroimage.2011.01.011].

72. H. Santosa, M. J. Hong, and K.-S. Hong, “Lateralization of music processing with noises in the auditory cortex: an fNIRS study,” Front. Behav. Neurosci. 8, Frontiers (2014) [doi:10.3389/fnbeh.2014.00418].

73. S. C. Herholz, A. R. Halpern, and R. J. Zatorre, “Neuronal Correlates of Perception, Imagery, and Memory for Familiar Tunes,” J. Cogn. Neurosci. 24(6), 1382–1397 (2012) [doi:10.1162/jocn_a_00216].

74. K. T. Kinder et al., “Systematic review of fNIRS studies reveals inconsistent chromophore data reporting practices,” Neurophotonics 9(4), 040601 (2022) [doi:10.1117/1.NPh.9.4.040601].

75. D. C. Van Essen et al., “The Human Connectome Project: a data acquisition perspective,” NeuroImage 62(4), 2222–2231 (2012) [doi:10.1016/j.neuroimage.2012.02.018].

76. M. A. Yücel et al., “fNIRS reproducibility varies with data quality, analysis pipelines, and researcher experience,” Commun. Biol. 8(1), 1149, Nature Publishing Group (2025) [doi:10.1038/s42003-025-08412-1].

