## Appendix for "All signals considered: Data quality partially explains inter-individual task differences in a large, open fNIRS dataset"

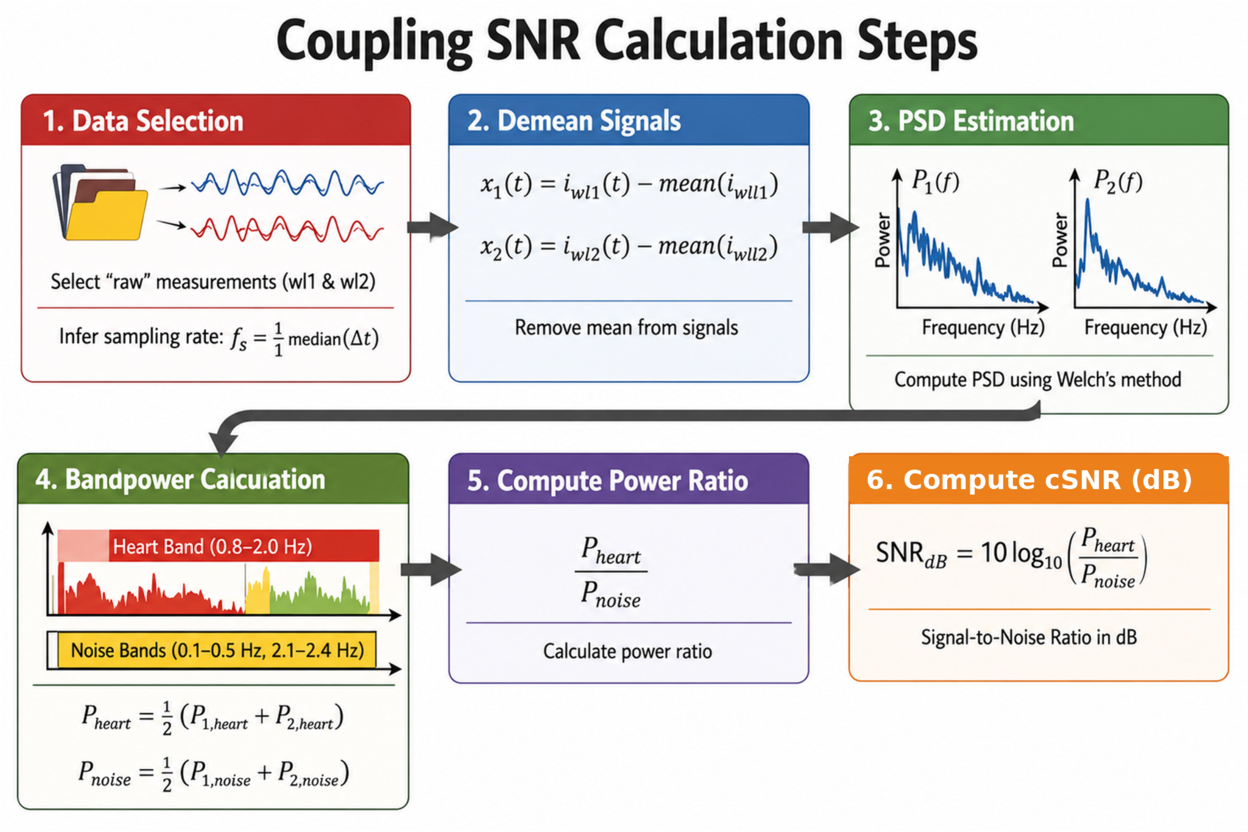


**Fig. S1** Overview of the calculation of the cSNR metric.

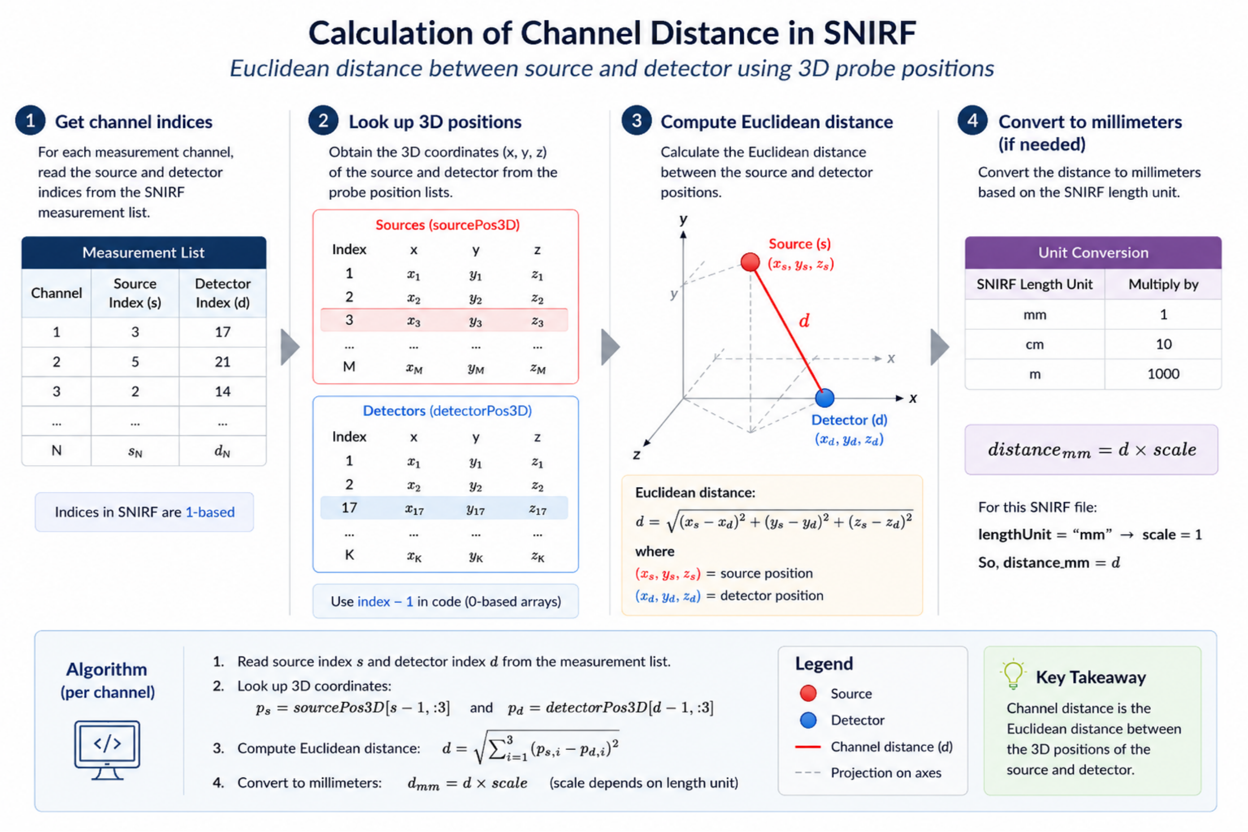


**Fig. S2** Overview of the process used to calculate the Euclidean source-detector distances from 3D probe positions.

**Table S1** Results of the ANCOVA investigating the effect of signal quality on inter-subject variability in motor task-induced activation across all participants (n = 57). Only channels with $p_{FDR} \leq .05$ are shown. Asterix denotes short-separation channels.

| **Quality Metric** | **Measure** | **Channel** | $\beta$ | **95% CI** | $F$**(1,55)** | $p$ | $p_{FDR}$ |
|---|---|---|---|---|---|---|---|
| cSNR | HbO | S10-D6 | 0.08 | [0.03, 0.12] | 11.65 | .001 | .041 |
| cSNR | HbO | S10-D7 | 0.07 | [0.03, 0.11] | 12.5 | .001 | .041 |
| cSNR | HbO | S25-D24 | 0.07 | [0.03, 0.11] | 11.89 | .001 | .041 |
| cSNR | HbO | S27-D26 | -0.06 | [-0.09, -0.03] | 13.17 | .001 | .041 |
| cSNR | HbR | S25-D24 | -0.09 | [-0.12, -0.05] | 18.79 | .000 | .008 |
| SCI | HbR | S10-D7 | -1.6 | [-2.50, -0.69] | 12.41 | .001 | .019 |
| SCI | HbR | S12-D40* | -1.87 | [-2.81, -0.93] | 15.91 | .000 | .013 |
| SCI | HbR | S17-D15 | -1.51 | [-2.33, -0.69] | 13.52 | .001 | .019 |
| SCI | HbR | S23-D51 | -1.58 | [-2.47, -0.69] | 12.66 | .001 | .019 |
| SCI | HbR | S25-D23 | -1.96 | [-2.82, -1.11] | 21.4 | .000 | .003 |
| SCI | HbR | S25-D53* | -1.54 | [-2.47, -0.61] | 11.11 | .002 | .030 |
| SCI | HbR | S27-D55* | -1.68 | [-2.60, -0.76] | 13.34 | .001 | .019 |
| SCI | Total | S12-D40* | -2.04 | [-3.12, -0.96] | 14.31 | .000 | .026 |
| SCI | Total | S24-D21 | 1.44 | [0.73, 2.14] | 16.78 | .000 | .019 |
| $CV_{\lambda=760}$ | HbR | S10-D7 | 0.09 | [0.04, 0.14] | 11.76 | .001 | .033 |
| $CV_{\lambda=760}$ | HbR | S12-D40* | 0.11 | [0.06, 0.16] | 19.88 | .000 | .005 |
| $CV_{\lambda=760}$ | HbR | S12-D7 | 0.08 | [0.03, 0.12] | 11.6 | .001 | .033 |
| $CV_{\lambda=760}$ | HbR | S17-D15 | 0.08 | [0.03, 0.12] | 11.04 | .002 | .035 |
| $CV_{\lambda=760}$ | HbR | S25-D23 | 0.1 | [0.05, 0.15] | 17.56 | .000 | .005 |
| $CV_{\lambda=760}$ | HbR | S25-D53* | 0.08 | [0.03, 0.14] | 10.74 | .002 | .035 |
| $CV_{\lambda=760}$ | HbR | S7-D35* | 0.08 | [0.03, 0.14] | 9.79 | .003 | .047 |
| $CV_{\lambda=760}$ | HbR | S9-D37* | 0.11 | [0.06, 0.16] | 17.22 | .000 | .005 |
| $CV_{\lambda=850}$ | HbO | S10-D7 | -0.13 | [-0.20, -0.06] | 13.98 | .000 | .020 |
| $CV_{\lambda=850}$ | HbO | S12-D7 | -0.12 | [-0.18, -0.06] | 16.99 | .000 | .017 |
| $CV_{\lambda=850}$ | HbO | S25-D23 | -0.14 | [-0.21, -0.07] | 15.25 | .000 | .017 |
| $SNR_{\lambda=760}$ | HbR | S12-D40* | -0.06 | [-0.10, -0.03] | 14.46 | .000 | .048 |
| $SNR_{\lambda=760}$ | HbR | S17-D15 | -0.05 | [-0.08, -0.02] | 12.7 | .001 | .048 |
| $SNR_{\lambda=760}$ | HbR | S25-D23 | -0.05 | [-0.09, -0.02] | 10.78 | .002 | .048 |
| $SNR_{\lambda=760}$ | HbR | S9-D23 | -0.06 | [-0.09, -0.02] | 10.86 | .002 | .048 |
| $SNR_{\lambda=760}$ | HbR | S9-D37* | -0.06 | [-0.10, -0.03] | 11.84 | .001 | .048 |
| $SNR_{\lambda=850}$ | HbO | S10-D6 | 0.09 | [0.05, 0.13] | 17.87 | .000 | .012 |
| $SNR_{\lambda=850}$ | HbO | S10-D7 | 0.07 | [0.03, 0.11] | 13.85 | .000 | .014 |
| $SNR_{\lambda=850}$ | HbO | S10-D8 | 0.07 | [0.03, 0.12] | 10.66 | .002 | .028 |
| $SNR_{\lambda=850}$ | HbO | S12-D7 | 0.07 | [0.03, 0.10] | 16.15 | .000 | .012 |
| $SNR_{\lambda=850}$ | HbO | S13-D8 | 0.06 | [0.02, 0.09] | 10.72 | .002 | .028 |
| $SNR_{\lambda=850}$ | HbO | S16-D11 | 0.06 | [0.02, 0.10] | 10.24 | .002 | .030 |
| $SNR_{\lambda=850}$ | HbO | S18-D15 | 0.07 | [0.02, 0.12] | 8.83 | .004 | .049 |
| $SNR_{\lambda=850}$ | HbO | S25-D23 | 0.08 | [0.04, 0.12] | 14.26 | .000 | .014 |
| $SNR_{\lambda=850}$ | HbO | S25-D24 | 0.07 | [0.03, 0.11] | 11.26 | .001 | .028 |
| $SNR_{\lambda=850}$ | HbO | S27-D23 | 0.07 | [0.03, 0.11] | 13.18 | .001 | .014 |
| $SNR_{\lambda=850}$ | HbO | S8-D6 | 0.07 | [0.02, 0.11] | 10.08 | .002 | .030 |
| $SNR_{\lambda=850}$ | HbO | S8-D8 | 0.08 | [0.03, 0.12] | 13.4 | .001 | .014 |

13

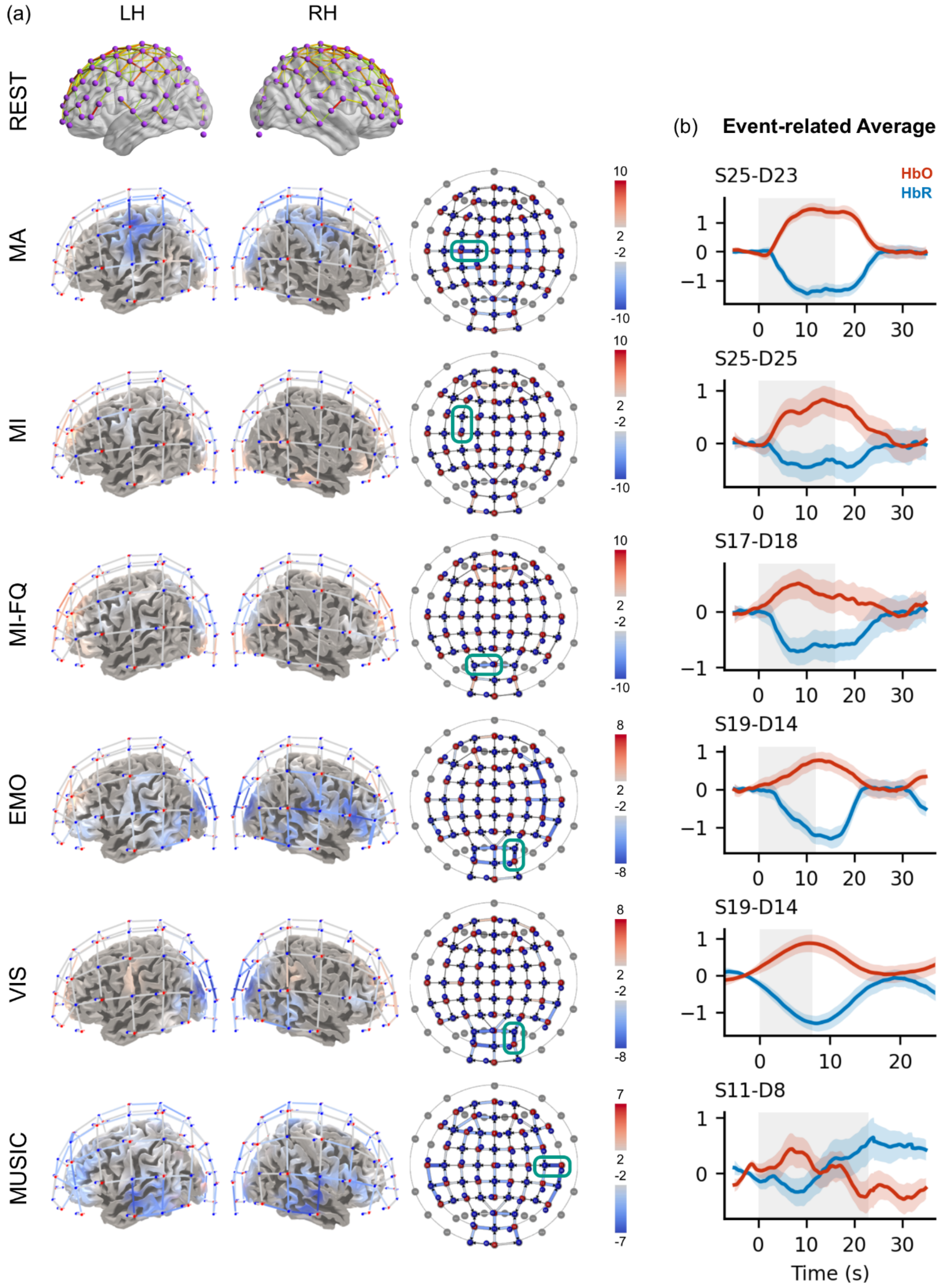

(a)
LH
RH
REST
MA
MI
MI-FQ
EMO
VIS
MUSIC
(b) Event-related Average
HbO
HbR
S25-D23
S25-D25
S17-D18
S19-D14
S19-D14
S11-D8
Time (s)

***Fig. S3*** (a) Connectivity map of the resting state task (top 5% strongest HbR connections at the group level) and RFX GLM results (t-maps) of the HbR data for each active task (n = 22), thresholded at $p \leq 0.05$. Results are displayed as exploratory surface projections on the LH = left hemisphere, RH = right hemisphere, and flat maps. For the visual, music and emotion task, all conditions are jointly contrasted with the baseline period. Channels with the lowest HbR t-value are highlighted on the flat maps in green for each task and the respective Event-related Averages of HbO and HbR are shown in (b). Grey shading indicates the task period. For the emotion task, the "Neutral-Happy" response is displayed. For the Visual Task, the "Threatening" category was chosen as an example. The music task shows the response to a "High-Valence, High-Arousal" stimulus. REST = Resting State, MA = Motor Action, MI = Motor Imagery, MI-FQ = Motor Imagery with frequency change, EMO = Emotion, VIS = Visual.
